# Mathematical Models of Retinitis Pigmentosa: The Trophic Factor Hypothesis

**DOI:** 10.1101/2021.03.25.436992

**Authors:** Paul A. Roberts

## Abstract

Retinitis pigmentosa (RP) is the term used to denote a group of inherited retinal-degenerative conditions that cause progressive sight loss. Individuals with this condition lose their light-sensitive photoreceptor cells, known as rods and cones, over a period of years to decades; degeneration starting in the retinal periphery, and spreading peripherally and centrally over time. RP is a rod-cone dystrophy, meaning that rod health and function are affected earlier and more severely than that of cones. Rods degenerate due to an underlying mutation, whereas the reasons for cone degeneration are unknown. A number of mechanisms have been proposed to explain secondary cone loss and the spatio-temporal patterns of retinal degeneration in RP. One of the most promising is the trophic factor hypothesis, which suggests that rods produce a factor necessary for cone survival, such that, when rods degenerate, cone degeneration follows. In this paper we formulate and analyse mathematical models of RP under the trophic factor hypothesis. These models are constructed as systems of reaction-diffusion partial differential equations in one spatial dimension, and are solved and analysed using a combination of numerical and analytical methods. We predict the conditions under which cones will degenerate following the loss of a patch of rods from the retina, the critical trophic factor treatment rate required to prevent cone degeneration following rod loss and the spatio-temporal patterns of cone loss that would result if the trophic factor mechanism alone were responsible for retinal degeneration.

## 1. Introduction

The group of inherited retinal diseases, known collectively as retinitis pigmentosa (RP), have been the subject of many decades of research. Despite this attention, many of the mechanisms underpinning RP have yet to be conclusively determined, though a number have been hypothesised (Hamel, 2006; Hartong et al., 2006). In this paper, we formulate mathematical models to describe and predict retinal degeneration in RP for one of the leading hypotheses: the trophic factor hypothesis.

The retina is the tissue layer at the back of the eye responsible for light detection (Fig. 1(a)). Retinal light-detecting cells, known as photoreceptors, come in two varieties: rods and cones. Rods confer peripheral, achromatic vision under low-light (scotopic) conditions, while cones confer high-acuity central, chromatic vision under well-lit (photopic) conditions (Oyster, 1999). Cones are mostly concentrated in the centre of the retina, known as the fovea, whereas rods dominate the mid- and far-peripheral retina (Curcio et al., 1990) (Fig. 1(b)). Both rods and cones are composed of inner and outer segments, together with an axon, ending in a synaptic terminal. Outer segments (OSs) contain a series of discs in which light-sensitive photopigments are embedded (Oyster, 1999). Each day, OS tips are shed and phagocytosed by the underlying retinal pigment epithelium (RPE), while new discs are generated where the OS meets the inner segment (Guérin et al., 1993; Jonnal et al., 2010, 2012; Kocaoglu et al., 2016; Pircher et al., 2011; Young, 1967, 1971, 1978; Young and Bok, 1969).

**Figure 1:**
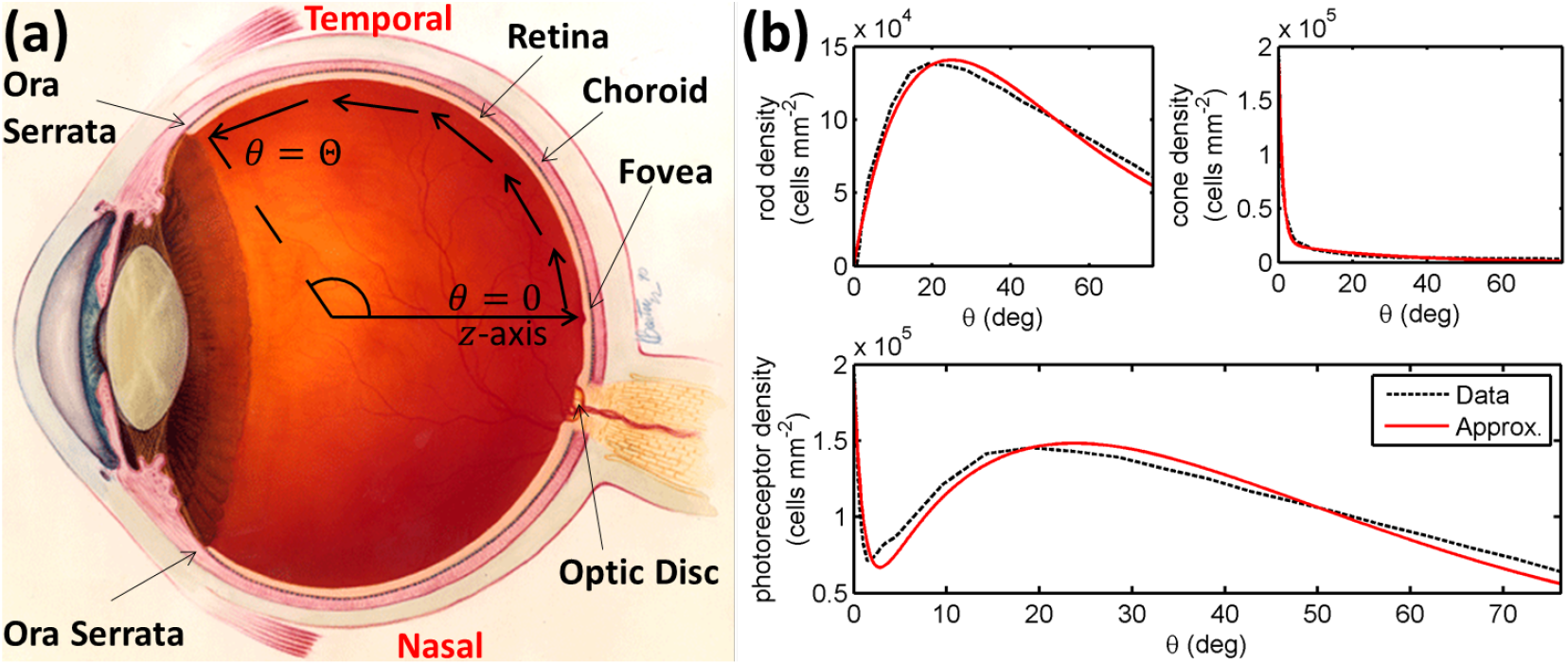
Diagrams of the human eye and retinal photoreceptor distribution (reproduced, with permission, from Roberts et al., 2017). (a) Diagram of the (right) human eye, viewed in the transverse plane, illustrating the model geometry. All models are posed on a domain spanning the region between the foveal centre, at *θ* = 0, and the ora serrata, at *θ* = Θ, along the temporal horizontal meridian, where *θ* measures the eccentricity. Figure originally reproduced, with modifications, from http://www.nei.nih.gov/health/coloboma/coloboma.asp, courtesy: National Eye Institute, National Institutes of Health (NEI/NIH). (b) Measured and fitted photoreceptor profiles, along the temporal horizontal meridian, in the human retina. Cone profile: 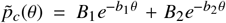, and rod profile: 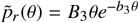 (see Table 1 for parameter values). The photoreceptor profile is the sum of the rod and cone profiles 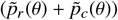. Experimental data provided by Curcio and published in Curcio et al. (1990).

RP is a rod-cone dystrophy, meaning that rod function and health are affected earlier and more severely than cone function and health (though there do exist rarer cone-rod dystrophy forms, and forms in which rods and cones are affected on a similar time scale, Hamel, 2006; Hartong et al., 2006). This is often because it is the rods, rather than the cones, which express the disease-causing mutation (though mutations can alternatively be expressed in the RPE, Daiger et al., 2007; Hamel, 2006; Hartong et al., 2006). Retinal degeneration (and hence visual field loss) typically initiates in the mid- or far-periphery of the retina, starting with small patches of photoreceptor loss (Cideciyan et al., 1998; García-Ayuso et al., 2013; Ji et al., 2012; Lee et al., 2011; Zhu et al., 2013) which spread and coalesce into larger regions, causing blind spots (scotomas). Degeneration spreads through the mid- and far-periphery until all peripheral vision is lost (tunnel vision), finally spreading to the central retina (macula), leading to the loss of central vision, and hence, total blindness (Grover et al., 1998; Hamel, 2006; Hartong et al., 2006). RP is the most common inherited retinal degeneration, with a prevalence of approximately 1 in 4000, and is presently untreatable (Hamel, 2006; Hartong et al., 2006; Musarella and MacDonald, 2011; Shintani et al., 2009).

Given that the disease-causing mutations are not typically expressed by cones, it is unclear what causes them to degenerate. Further, the reasons behind the spatio-temporal patterns of retinal degeneration in general, and for which any particular degenerate patch expands or remains stable, remain a mystery (Hamel, 2006; Hartong et al., 2006). A number of mechanisms have been proposed to explain these phenomena, most notably trophic factor depletion (Aït-Ali et al., 2015; Léveillard et al., 2004; Mei et al., 2016), oxygen toxicity (Stone et al., 1999; Travis et al., 1991; Valter et al., 1998), metabolic dysregulation (Punzo et al., 2009, 2012), toxic substances (Ripps, 2002) and microglia (Gupta et al., 2003). In this paper, we consider the first of these, often denoted as the trophic factor hypothesis. This hypothesis suggests that rods produce a trophic factor necessary for cone survival, such that, when rods degenerate, the trophic factor is depleted, and cone degeneration follows. Such a factor, known as rod-derived cone viability factor (RdCVF), has been chemically identified by Léveillard et al. (2004), and shown to slow cone degeneration and preserve cone function in chick, rat and mouse models (Fintz et al., 2003; Léveillard et al., 2004; Mohand-Saïd et al., 1998, 2000, 1997; Yang et al., 2009). RdCVF promotes cone survival by binding to the photoreceptor transmembrane protein Basigin-1 (BSG1), which in turn binds to the glucose transporter GLUT1, increasing cone glucose uptake and stimulating aerobic glycolysis (Aït-Ali et al., 2015). For reviews, see Clérin et al. (2020); Léveillard and Aït-Ali (2017); Léveillard and Sahel (2010, 2017); Mohand-Said et al. (2001).

In this paper, we construct mathematical models to describe and predict retinal degeneration and preservation under the trophic factor hypothesis. As such, we are isolating the trophic factor mechanism, in a manner that would be almost impossible to achieve experimentally, to investigate what would happen if this mechanism were solely responsible for cone degeneration in RP. Our models are formulated as systems of reaction-diffusion partial differential equations (PDEs) in one spatial dimension, and solved and analysed using a combination of numerical and analytical techniques. We use these models to address three key questions: (i) under what conditions will the loss of a patch of rods lead to cone degeneration?, (ii) what is the critical trophic factor treatment dose required to prevent cone degeneration following the loss of a patch of rods?, and (iii) what spatio-temporal patterns of cone degeneration can be explained via the trophic factor mechanism alone?

In earlier work, we developed the first and only models to consider the oxygen toxicity hypothesis, and to take the key step of accounting for the distribution of photoreceptors (Roberts et al., 2017, 2018, see also, Roberts et al. 2016b). These models are formulated as systems of reaction-diffusion PDEs in one and two spatial dimensions, and were used to answer the long-standing question as to the spatio-temporal patterns of retinal degeneration that would result from this mechanism alone. These studies further predicted the propagation speed of wavefronts of hyperoxia-driven retinal degeneration, the conditions under which degenerate patches will expand or remain stable, and the effects of treatment with antioxidants and trophic factors.

Burns et al. (2002) developed the only (1D PDE) model to consider the toxic substance hypothesis. This model captures the patchy loss of photoreceptors seen in the early stages of RP and, under the right conditions, replicates the exponential decline in photoreceptors observed by Clarke et al. (2000, 2001).

Lastly, Camacho *et al.* have produced a series of ordinary differential equation models of the trophic factor hypothesis, which focus on the biochemical details, rather than the spatial spread, of retinal degeneration (Colón Vélez et al., 2003; Camacho et al., 2010; Camacho and Wirkus, 2013; Camacho et al., 2014, 2016a,b,c, 2019, 2020, 2021; Wifvat et al., 2021). Their models replicate the rhythmic shedding and renewal of photoreceptor OSs; predict possible disease stages through which a retina may pass towards total blindness, recapitulating known RP phenotypes; identify key reactions and processes; and predict optimum treatment strategies. The work presented here is complementary to Camacho *et al.*’s models, focusing on the spatial spread of degeneration, while remaining relatively agnostic as to the underlying biochemical details (and hence is compatible with a range of trophic factor-related biochemical mechanisms). For further details on the mathematical modelling of the retina, including RP, see our review: Roberts et al. (2016a).

The models developed and analysed in this paper are the first trophic factor hypothesis models to be spatially-resolved, or to account for photoreceptor distribution. As such, these are the first trophic factor models capable of answering the three key questions posed above (though we note that Camacho *et al.*’s work considers spatially-uniform analogues of questions (i) and (ii), see especially, Camacho et al., 2014, 2020).

The remainder of this paper is structured as follows. In Section 2, we formulate and non-dimensionalise our mathematical models. In Section 3, we consider the steady-state model, performing asymptotic analyses (Section 3.1), and comparing numerical and analytical solutions (Section 3.2) in the untreated (Section 3.2.1) and treated (Section 3.2.2) cases. In Section 4, we consider numerical solutions to the full dynamic (time dependent) model, in both the untreated (Section 4.1) and treated (Section 4.2) cases. Lastly, in Section 5, we discuss our results and suggest directions for future research.

## 2. Model formulation

We begin by formulating a model to describe the photoreceptor and trophic factor dynamics within the human retina. Given that the retina lines the essentially spherical inner surface of the eye, we pose our model in spherical polar coordinates (*r,θ,ϕ*), with origin at the centre of the vitreous body and orientated such that the z-axis passes outwards through the centre of the fovea (see Fig. 1(a)). We make two geometrically simplifying assumptions. First, we assume that the retina is symmetric about the *z*-axis, such that the azimuthal dimension, *ϕ* (rad), can be neglected. While this ignores the optic disc (from which photoreceptors are absent) and other inhomogeneities in rod and cone densities, this is a reasonable simplification to make in the first spatial model of this disease mechanism. Second, since the retina’s width (~80–320 *μ*m, Webvision, https://webvision.med.utah.edu/) is approximately two orders of magnitude smaller than its radius of curvature (~1.2 cm, Oyster, 1999), we depth-average through the retina, assuming a fixed radius *r* = *R* (m). Thus, we have reduced the geometry to (*R, θ*); posing our model on the one-dimensional domain *θ* ∈ [0, Θ] (rad), where *θ* = 0 (rad) is located at the foveal centre and *θ* = Θ (rad) is located at the ora serrata.

We consider two scenarios, leading to alternate forms of the governing equations:

- Scenario 1: without (w/o) cone regeneration;
- Scenario 2: with cone regeneration.

In Scenario 1, we model the cone photoreceptor density, *p_c_*(*θ, t*) (photoreceptors m^−2^), with its associated initial value, *p_c_nit__*(*θ*) (photoreceptors m^−2^). In this case, cone degeneration corresponds to the loss of whole cone photoreceptor cells. Thus, given that photoreceptors cannot be regrown once they are lost, recovery is impossible. In Scenario 2, we model the local cone OS length, 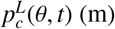, with its associated initial value, 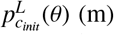. In this case, cone degeneration corresponds to the shortening of cone OS, while leaving the rest of the photoreceptor intact, such that cone density remains fixed at *p_c_init__* (*θ*) (photoreceptors m^−2^) for all time. As such, cone regeneration is possible in the form of OS regrowth (see Guérin et al., 1993, and also Chrysostomou et al., 2008; Lee et al., 2012). Since prolonged exposure to adverse conditions will lead to the eventual loss of the whole photoreceptor, it is assumed in this scenario that adverse conditions are either mild or relatively brief.

We construct a system of partial differential equations (PDEs) in terms of the dependent variables: trophic factor concentration, *f*(*θ, t*) (M), rod photoreceptor density, *p_r_*(*θ, t*) (photoreceptors m^−2^), and cone photoreceptor density, *p_c_*(*θ, t*) (photoreceptors m^−2^), or local cone OS length, 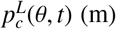; as functions of the independent variables: polar angle, *θ* (rad), and time, *t* (s), where *t* > 0 (s).

The trophic factor equation takes the following form,

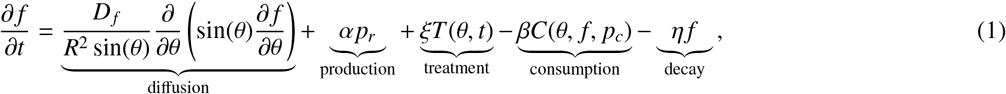

where *R* (m), the radial position of the retina, *D_f_* (m^2^s^−1^), the trophic factor diffusivity, *α* (Mm^2^photoreceptors^−1^ s^−1^), the rate of trophic factor production by rods, *ξ* (M s^−1^), the rate of supply of trophic factor through treatment, *β* (m^2^photoreceptors^−1^s^−1^), the rate of trophic factor consumption by cones, and *η* (s^−1^), the rate of trophic factor decay, are positive constants. The treatment function is defined as follows

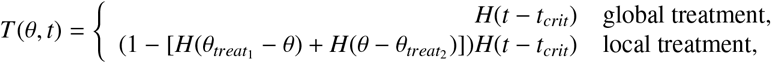

where trophic factor (RdCVF) treatment is given from time *t* = *t_crit_* > 0 (s), and can be applied either globally, across the whole domain *θ* ∈ [0, Θ] (rad), or locally, within the region *θ* ∈ (*θ*_*treat*_1__, *θ*_*treat*_2__) (rad) only. We define the Heaviside step function, *H*(·), such that

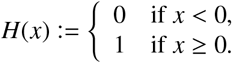

The trophic factor consumption function is defined as follows

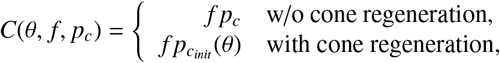

where *p_c_init__*(*θ*) (photoreceptors m^−2^), the initial cone density, is defined below. In Scenario 1, cone density may decrease, such that the rate of TF consumption changes both with decreasing *p_c_*(*θ, t*) and changing *f* (*θ, t*). By contrast, in Scenario 2, the cone density remains fixed, such that the rate of TF consumption changes with *f* (*θ, t*) only.

The rod equation is as follows

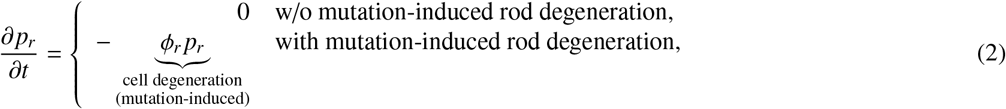

where *ϕ_r_* (s^−1^), the rate of mutation-induced rod degeneration, is a positive constant. We keep the rod density constant in the case where we seek to focus on the photoreceptor dynamics following the formation of a degenerate patch (from which rods and/or cones are absent), while an exponential decline in rods more faithfully represents the natural disease progression. Therefore, in the case without mutation-induced rod degeneration, the rod density remains at its initial value, *p_r_*(*θ, t*) = *p_r_init__* (*θ*) (photoreceptors m^−2^), while in the case with mutation-induced rod degeneration, *p_r_*(*θ, t*) = *p_r_init__*(*θ*)*e^−ϕ_r_t^* (photoreceptors m^−2^), provided there is no delay in onset or interruption of degeneration.

In Scenario 1, the cone photoreceptor dynamics are given by

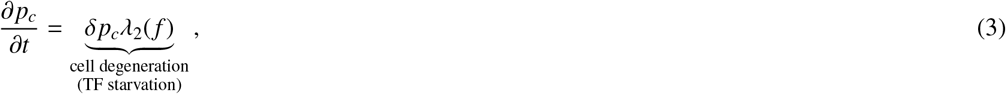

where *δ* (s ^−1^), the rate of trophic factor (TF) starvation-induced cone degeneration, is a positive constant. In Scenario 2, the cone photoreceptor dynamics are given by

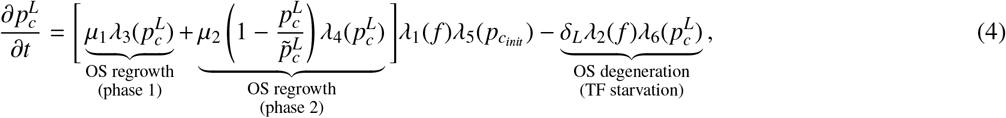

where 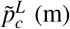, the healthy cone OS length, *μ*_1_ (s^−1^), the growth rate of cone OS in phase 1, *μ*_2_ (s^−1^), the growth rate of cone OS in phase 2, and *δ_L_* (ms^−1^), the rate of trophic factor starvation-induced cone OS degeneration, are positive constants. In Scenario 1, cone cells degenerate exponentially, while in Scenario 2, individual cone OS degenerate at a constant rate.

Model fitting to cone OS regrowth data from Guérin et al. (1993) reveals that cone OS regrowth is well-described by a two-phase model. Phase 1, constant growth, occurs for cone OS lengths between 0 and 0.33 as a proportion of their full length. In this early, rapid regrowth phase, new OS discs are regenerated, but OS shedding does not occur. Phase 2, hyperbolic growth, occurs for cone OS lengths between 0.33 and 1 as a proportion of their full length. In this later, slower regrowth phase, new OS discs continue to be regenerated, while OS shedding is re-established. The functions *λ*_1–6_ are given by

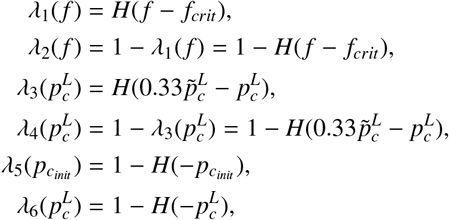

where *f_crit_* (M), the trophic factor threshold concentration, is a positive constant.

The functions *λ*_1_(*f*) and *λ*_2_(*f*) determine when the cone regrowth and degeneration terms are active. Cone degeneration occurs when *f* < *f_crit_* (TF starvation) in both Scenarios 1 and 2, while cone (OS) regrowth occurs when *f* ≥ *f_crit_* in Scenario 2, where TF levels are sufficient to maintain cones in health. The functions 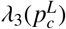 and 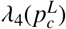 determine whether phase 1 or phase 2 cone OS regrowth is active. Phase 1 is active when 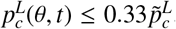, while phase 2 is active when 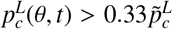. The function *λ*_5_(*p_c_init__*) prevents cone OS regrowth in regions from which cones are absent initially. The function 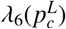 halts cone OS degeneration when their length reaches zero, preventing cone OS length becoming negative.

Lastly, we close the system by imposing boundary and initial conditions. We impose zero-flux boundary conditions at both the fovea (*θ* = 0) and ora serrata (*θ* = Θ):

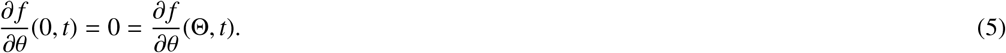

A zero-flux condition is required by symmetry at *θ* = 0, while the zero-flux condition at *θ* = Θ is justified by the assumption that TF cannot escape from the retina at the ora serrata.

We impose the following initial conditions:

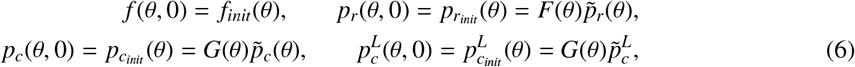

where *f_init_*(*θ*) (M) is the steady-state solution to Eqs. (1) and (5) with *p_r_* = *p_r_init__*(*θ*) (photoreceptors m^−2^), *p_c_* = *p_c_init__*(*θ*) (photoreceptors m^−2^) and *T*(*θ*, 0) = 0 (no treatment), while 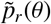 (photoreceptors m^−2^), the healthy rod distribution, and 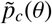 (photoreceptors m^−2^), the healthy cone distribution are defined as

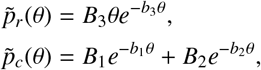

where *B*_1_ (photoreceptors m^−2^), *B*_2_ (photoreceptors m^−2^), *B*_3_ (photoreceptors m^−2^ rad^−1^), *b*_1_ (rad^−1^), *b*_2_ (rad^−1^) and *b*_3_ (rad^−1^) are positive constants, obtained by fitting to physiological data (see Fig. 1(b), Fig. 2(a) and Curcio et al., 1990). We make the simplifying assumption that healthy cone OS length, 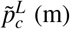, is constant across the retina. We consider two types of initial conditions for rods and cones: healthy and containing a degenerate patch. Which of these situations obtain is determined by the functions *F*(*θ*) and *G*(*θ*), which are given by

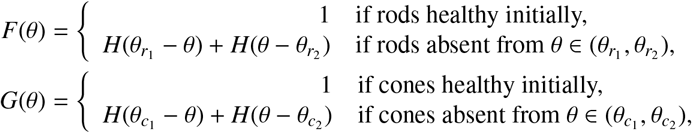

where *θ*_*r*_1__ (rad) and *θ*_*r*_2__ (rad) are the left- (central) and right-hand (peripheral) boundaries of a patch of rod degeneration, and *θ*_*c*_1__ (rad) and *θ*_*c*_2__ (rad) are the left- (central) and right-hand (peripheral) boundaries of a patch of cone degeneration. Example initial conditions are presented in Fig. 2(b)–(d).

**Figure 2:**
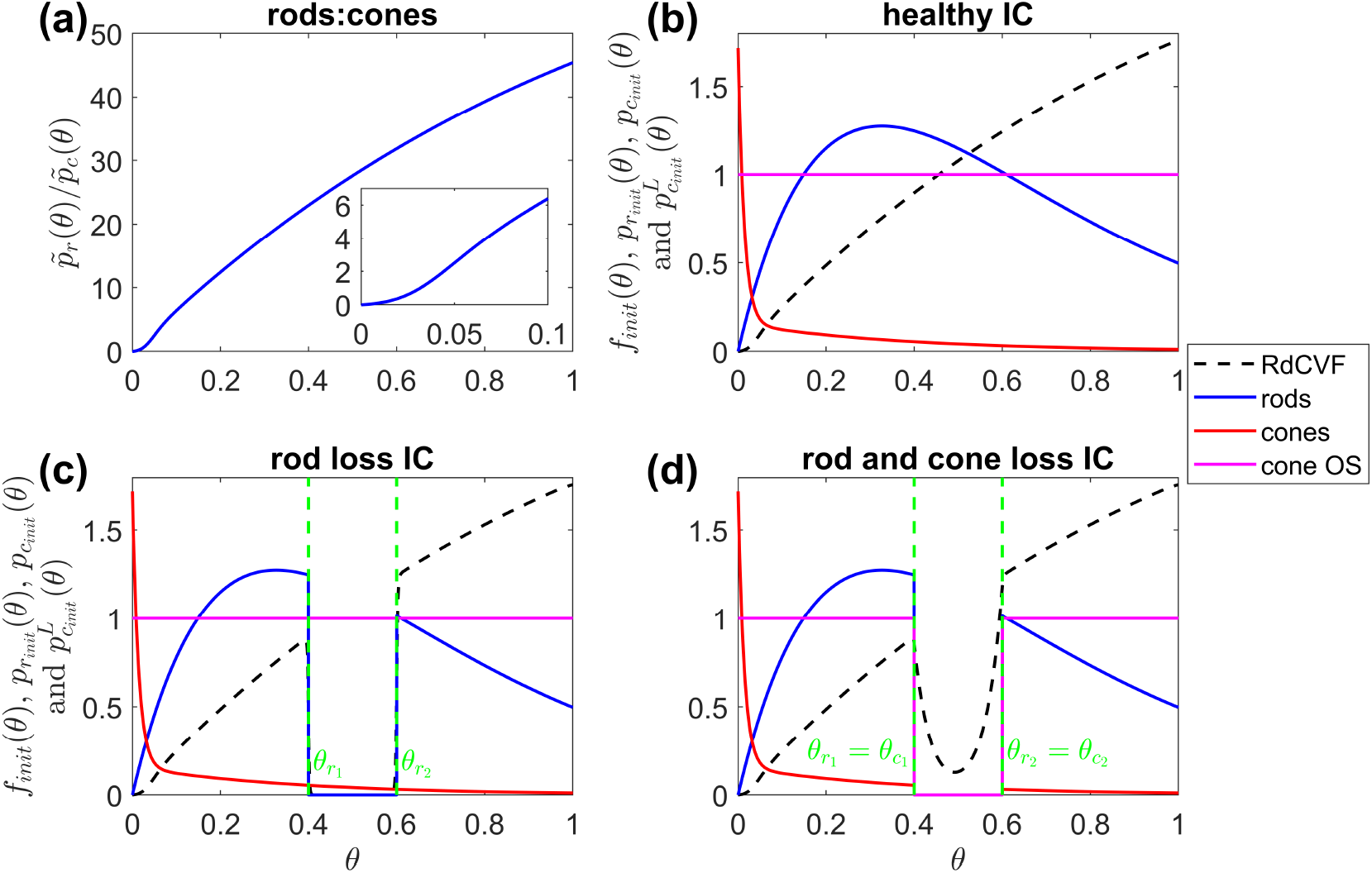
Ratio of rods to cones and exemplar initial conditions. (a) variation in the healthy rod:cone ratio, 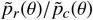, with eccentricity (inset shows magnified bottom-left corner). (b)–(d) examples of typical initial conditions (ICs) in the healthy retina (b), with a patch of rod loss (c), and with a patch of rod and cone loss (d); the legend applies to (b)–(d) only. The green dashed vertical lines mark the boundaries of rod (and cone) loss. To obtain *f_init_*(*θ*) in (b)–(d), Eqs. (7) and (11) were solved using the finite difference method, with 4001 mesh points, where 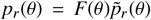 and 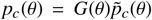, and without treatment. (b) *F*(*θ*) = 1 and *G*(*θ*) = 1; (c) *F*(*θ*) = *H*(*θ*_*r*_1__ – *θ*) + *H*(*θ* – *θ*_*r*_2__) and *G*(*θ*) = 1; (d) *F*(*θ*) = *H*(*θ*_*r*_1__ – *θ*) + *H*(*θ* < *θ*_*r*_2__) and *G*(*θ*) = *H*(*θ*_*c*_1__ – *θ*) + *H*(*θ* – *θ*_*c*_2__). Parameter values: *ξ* = 0; (*θ*_*r*_1__, *θ*_*r*_2__) = (0.4,0.6) in (c) and (d); (*θ*_*c*_1__, *θ*_*c*_2__) = (0.4,0.6) in (d). Remaining parameter values as in Table 2.

Parameter values associated with the dimensional model can be found in Table 1. See the Supplementary Materials for justification of parameter values.

**Table 1:**
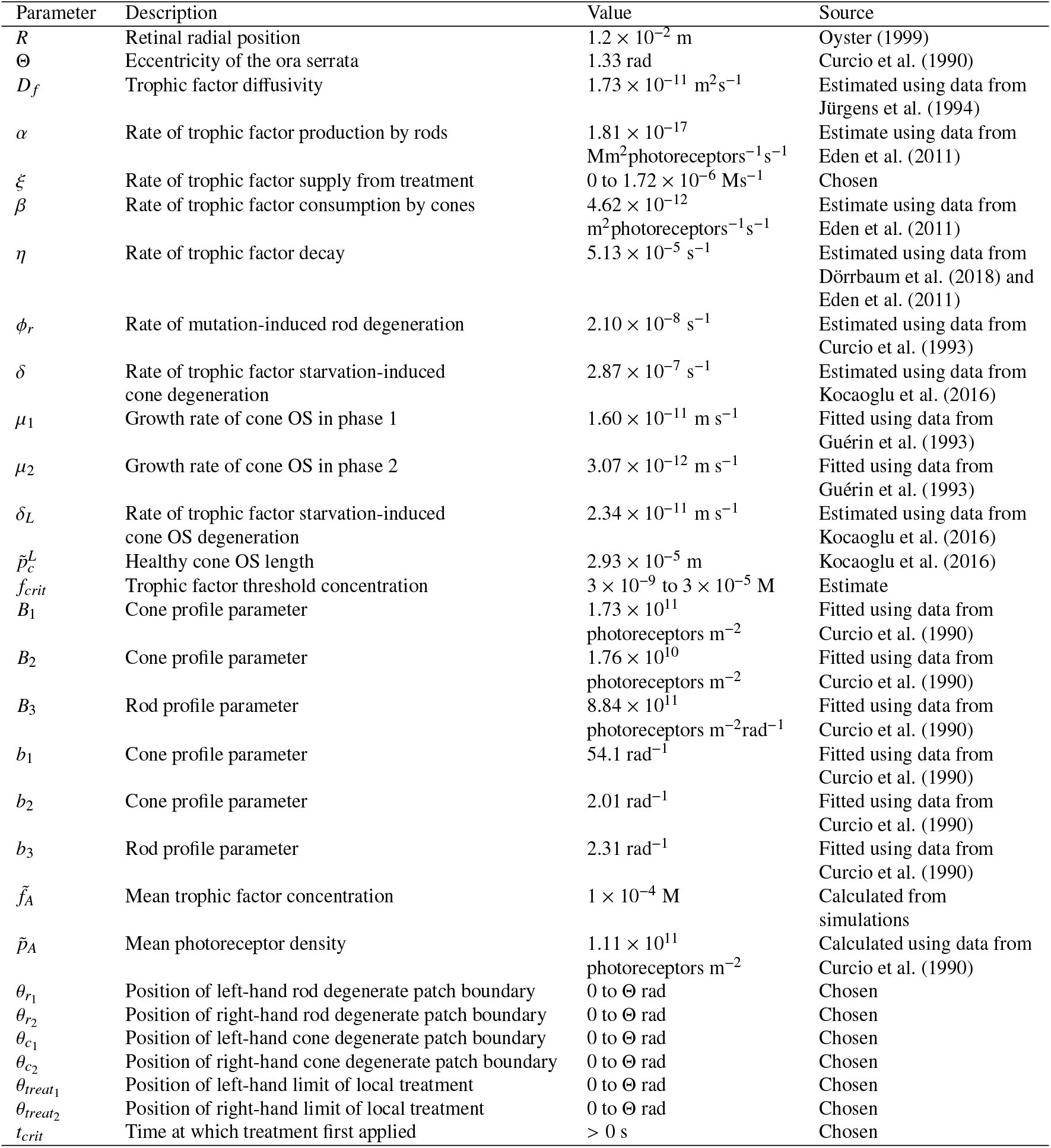
Parameters employed in the dimensional model (Eqs. (1)–(6)). Parameters for which only the source is given are taken directly from the literature, those described as ‘calculated’ or ‘fitted’ are computed using data from the literature or from simulations, those described as ‘estimated’ are either estimated from relevant data in the literature or on the basis of biological reasoning, and those described as ‘chosen’ are free for us to choose. Values are given to three significant figures. See Supplementary Materials for justification of parameter values.

| Parameter | Description | Value | Source |
| --- | --- | --- | --- |
| $R$ | Retinal radial position | $1.2 \times 10^{-2}$ m | Oyster (1999) |
| $\Theta$ | Eccentricity of the ora serrata | 1.33 rad | Curcio et al. (1990) |
| $D_f$ | Trophic factor diffusivity | $1.73 \times 10^{-11}$ m<sup>2</sup>s<sup>-1</sup> | Estimated using data from Jürgens et al. (1994) |
| $\alpha$ | Rate of trophic factor production by rods | $1.81 \times 10^{-17}$ Mm<sup>2</sup>photoreceptors<sup>-1</sup>s<sup>-1</sup> | Estimate using data from Eden et al. (2011) |
| $\xi$ | Rate of trophic factor supply from treatment | 0 to $1.72 \times 10^{-6}$ Ms<sup>-1</sup> | Chosen |
| $\beta$ | Rate of trophic factor consumption by cones | $4.62 \times 10^{-12}$ m<sup>2</sup>photoreceptors<sup>-1</sup>s<sup>-1</sup> | Estimate using data from Eden et al. (2011) |
| $\eta$ | Rate of trophic factor decay | $5.13 \times 10^{-5}$ s<sup>-1</sup> | Estimated using data from Dörrbaum et al. (2018) and Eden et al. (2011) |
| $\phi_r$ | Rate of mutation-induced rod degeneration | $2.10 \times 10^{-8}$ s<sup>-1</sup> | Estimated using data from Curcio et al. (1993) |
| $\delta$ | Rate of trophic factor starvation-induced cone degeneration | $2.87 \times 10^{-7}$ s<sup>-1</sup> | Estimated using data from Kocaoglu et al. (2016) |
| $\mu_1$ | Growth rate of cone OS in phase 1 | $1.60 \times 10^{-11}$ m s<sup>-1</sup> | Fitted using data from Guérin et al. (1993) |
| $\mu_2$ | Growth rate of cone OS in phase 2 | $3.07 \times 10^{-12}$ m s<sup>-1</sup> | Fitted using data from Guérin et al. (1993) |
| $\delta_L$ | Rate of trophic factor starvation-induced cone OS degeneration | $2.34 \times 10^{-11}$ m s<sup>-1</sup> | Estimated using data from Kocaoglu et al. (2016) |
| $\tilde{p}_c^L$ | Healthy cone OS length | $2.93 \times 10^{-5}$ m | Kocaoglu et al. (2016) |
| $f_{crit}$ | Trophic factor threshold concentration | $3 \times 10^{-9}$ to $3 \times 10^{-5}$ M | Estimate |
| $B_1$ | Cone profile parameter | $1.73 \times 10^{11}$ photoreceptors m<sup>-2</sup> | Fitted using data from Curcio et al. (1990) |
| $B_2$ | Cone profile parameter | $1.76 \times 10^{10}$ photoreceptors m<sup>-2</sup> | Fitted using data from Curcio et al. (1990) |
| $B_3$ | Rod profile parameter | $8.84 \times 10^{11}$ photoreceptors m<sup>-2</sup>rad<sup>-1</sup> | Fitted using data from Curcio et al. (1990) |
| $b_1$ | Cone profile parameter | $54.1$ rad<sup>-1</sup> | Fitted using data from Curcio et al. (1990) |
| $b_2$ | Cone profile parameter | $2.01$ rad<sup>-1</sup> | Fitted using data from Curcio et al. (1990) |
| $b_3$ | Rod profile parameter | $2.31$ rad<sup>-1</sup> | Fitted using data from Curcio et al. (1990) |
| $\tilde{f}_A$ | Mean trophic factor concentration | $1 \times 10^{-4}$ M | Calculated from simulations |
| $\tilde{p}_A$ | Mean photoreceptor density | $1.11 \times 10^{11}$ photoreceptors m<sup>-2</sup> | Calculated using data from Curcio et al. (1990) |
| $\theta_{r1}$ | Position of left-hand rod degenerate patch boundary | 0 to $\Theta$ rad | Chosen |
| $\theta_{r2}$ | Position of right-hand rod degenerate patch boundary | 0 to $\Theta$ rad | Chosen |
| $\theta_{c1}$ | Position of left-hand cone degenerate patch boundary | 0 to $\Theta$ rad | Chosen |
| $\theta_{c2}$ | Position of right-hand cone degenerate patch boundary | 0 to $\Theta$ rad | Chosen |
| $\theta_{treat1}$ | Position of left-hand limit of local treatment | 0 to $\Theta$ rad | Chosen |
| $\theta_{treat2}$ | Position of right-hand limit of local treatment | 0 to $\Theta$ rad | Chosen |
| $t_{crit}$ | Time at which treatment first applied | > 0 s | Chosen |

### 2.1. Non-dimensionalisation

We recast the dimensional model in non-dimensional form to identify the dominant terms and simplify the equations. We scale the independent and dependent variables, and initial conditions as follows:

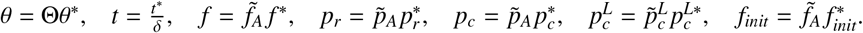

Consequently, the scaled system is posed on the domain *θ*\* ∈ [0,1]. We rescale time by the rate of cone degeneration, *δ*, since this is the timescale of interest. The scaling factor 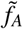 is the mean trophic factor concentration under healthy conditions, found by taking the mean of the solution to Eqs. (1) and (5) at steady-state, with 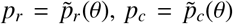 and *T*(*θ*) = 0 (no treatment). The scaling factor 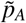 is the mean total photoreceptor density (rods and cones) under healthy conditions.

We define the following dimensionless parameters:

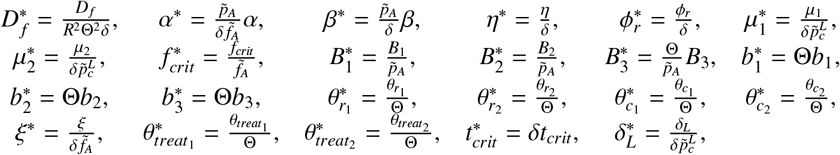

and note that *T*(*θ, t*) = *T*\*(*θ*, t*\*), where 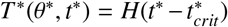 or 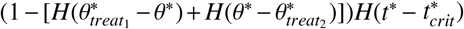 (in the global and local treatment cases respectively); 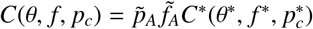, where 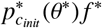 or 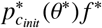 (in the without cone regeneration and with cone regeneration cases respectively); 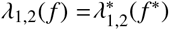, where 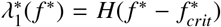 and 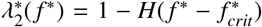; 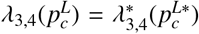, where 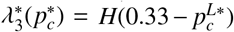 and 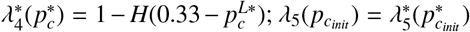 where 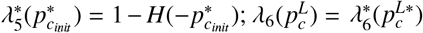, where 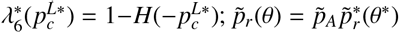, where 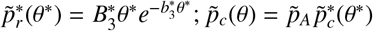, where 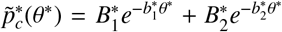; *F*(*θ*) = *F*\*(*θ*\*), where *F*\*(*θ*\*) = 1 or 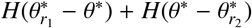 (in the healthy and degenerate patch cases respectively); and *G*(*θ*) = *G*\*(*θ*\*), where *G*\*(*θ*\*) = 1 or 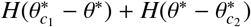 (in the healthy and degenerate patch cases respectively).

Applying the above scalings to the dimensional governing equations, boundary and initial conditions (Eqs. (1)–(6)) and dropping the stars, we obtain the following dimensionless governing equations

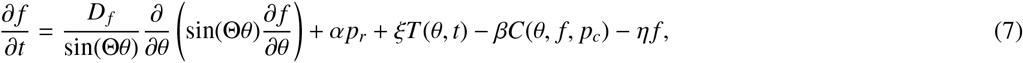

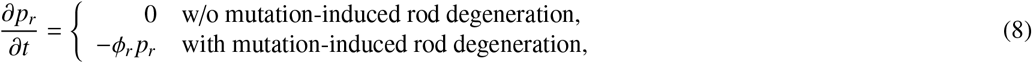

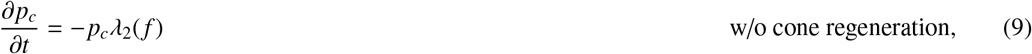

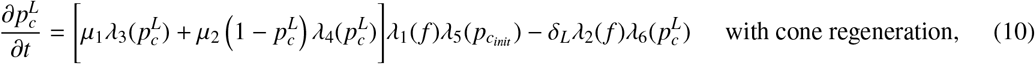

boundary conditions,

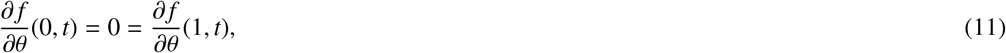

and initial conditions,

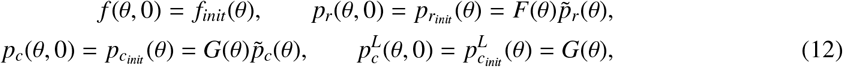

where *f_init_*(*θ*) is the steady-state solution to Eqs. (7) and (11) with *p_r_* = *p_r_init__*(*θ*), *p_c_* = *p_c_init__*(*θ*) and *T*(*θ*) = 0 (no treatment). See Table 2 for the dimensionless parameter values.

**Table 2:** Parameters employed in the non-dimensional model (Eqs. (7)–(12)). Values are given to three significant figures.

| Parameter | Description | Value |
| --- | --- | --- |
| $D_f$ | Trophic factor diffusivity | 0.237 |
| $\alpha$ | Rate of trophic factor production by rods | $7.01 \times 10^4$ |
| $\xi$ | Rate of trophic factor supply from treatment | 0 to $6 \times 10^4$ |
| $\beta$ | Rate of trophic factor consumption by cones | $1.79 \times 10^6$ |
| $\eta$ | Rate of trophic factor decay | $1.79 \times 10^2$ |
| $\phi_r$ | Rate of mutation-induced rod degeneration | $7.33 \times 10^{-2}$ |
| $\mu_1$ | Growth rate of cone OS in phase 1 | 1.90 |
| $\mu_2$ | Growth rate of cone OS in phase 2 | 0.365 |
| $\delta_L$ | Rate of trophic factor starvation-induced cone OS degeneration | 2.79 |
| $f_{crit}$ | Trophic factor threshold concentration | $3 \times 10^{-5}$ or 0.3 |
| $B_1$ | Cone profile parameter | 1.56 |
| $B_2$ | Cone profile parameter | 0.158 |
| $B_3$ | Rod profile parameter | 10.6 |
| $b_1$ | Cone profile parameter | 71.8 |
| $b_2$ | Cone profile parameter | 2.67 |
| $b_3$ | Rod profile parameter | 3.06 |
| $\theta_{r1}$ | Position of left-hand rod degenerate patch boundary | 0 to 1 |
| $\theta_{r2}$ | Position of right-hand rod degenerate patch boundary | 0 to 1 |
| $\theta_{c1}$ | Position of left-hand cone degenerate patch boundary | 0 to 1 |
| $\theta_{c2}$ | Position of right-hand cone degenerate patch boundary | 0 to 1 |
| $\theta_{treat1}$ | Position of left-hand limit of local treatment | 0 to 1 |
| $\theta_{treat2}$ | Position of right-hand limit of local treatment | 0 to 1 |
| $t_{crit}$ | Time at which treatment first applied | $> 0$ |

## 3. Results: steady-state problem

In this section we consider the RdCVF Eqs. (7) and (11) at steady-state. Using a combination of asymptotic analysis and numerical simulations we seek to determine the spatial extent of a patch of cone loss, consequent upon the loss of a patch of rods, and the critical RdCVF treatment rate required to prevent cone degeneration following a patch of rod loss.

### 3.1. Asymptotic analyses

There are a variety of scenarios to be considered:

- Rod loss only / rod and cone loss: either rods alone are removed from a patch, or rods and cones are removed from a patch, providing upper and lower bounds, respectively, on the predicted spatial extent of cone loss and critical treatment rate (see Section 3.2.1 for further details);
- Wide/narrow patch of photoreceptor loss: wide patches must be treated in an asymptotically distinct way from narrow patches (see below), and represent late and early disease stages respectively;
- No treatment / treatment: in the untreated case we are interested in the extent of cone loss and in the treatment case we are interested in the critical RdCVF treatment rate required to prevent cone loss;

– Global/local treatment: RdCVF treatment may be applied globally, across the whole retina, or locally, within the degenerate rod patch only. We are interested to see to what extent the critical treatment rate varies with global vs. local application.

We consider 8 cases (of a possible 12):

1. Wide patch of rod loss without treatment;
2. Narrow patch of rod loss without treatment;
3. Wide patch of rod and cone loss without treatment;
4. Narrow patch of rod and cone loss without treatment;
5. Wide patch of rod loss with global treatment;
6. Wide patch of rod loss with local treatment;
7. Narrow patch of rod loss with global treatment;
8. Narrow patch of rod loss with local treatment.

We do not consider rod and cone loss with treatment (4 cases) since we are only interested in calculating the upper bound on the critical treatment rate.

In the next section (3.1.1) we will use the following variables and parameters, defined here for ease of reference:

- 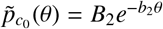;
- 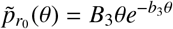;
- 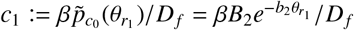;
- 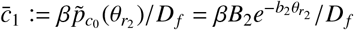;
- 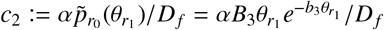;
- 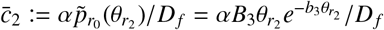.

### 3.1.1. Wide patch of rod loss without treatment

In this section we perform an asymptotic analysis with the aim of deriving analytical expressions for the boundaries of cone loss following the formation of a degenerate rod patch. We assume that cone loss will occur in the region for which the trophic factor concentration *f* < *f_crit_* local to the degenerate rod patch. Thus, the boundaries of the degenerate cone patch will be given as the eccentricities at which *f* = *f_crit_*. To determine where this occurs, we seek a leading order solution for *f*(*θ*) to Eqs. (7) and (11) at steady-state, for prescribed 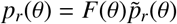 and 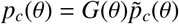, where *F*(*θ*) = *H*(*θ*_*r*_1__ – *θ*) + *H*(*θ* − *θ*_*r*_2__) and *G*(*θ*) = 1.

We choose *ϵ* = *O*(10^−2^), noting that we set *ϵ* = 10^−2^ for all plots, scaling parameters as follows: *α* = *ϵ*^−2^*α*′, *β* = *ϵ*^−3^*β*′, *η* = *ϵ*^−1^*η*′ and 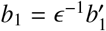 (using the standard parameter values as in Table 2). All remaining parameters are assumed to be *O*(1). We note that we could have chosen values for *α* and *β* two orders of magnitude smaller than their present values, resulting in an alternative distinguished limit. This alternative scaling will be explored in a future publication. We also scale the dependent variable 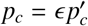, and assume *f* = *O*(1) and *p_r_* = *O*(1). This scaling on the dependent variables is valid in the region *θ* ∈ (*ϵ*, 1] here and in all rod loss only cases. For completeness, we note that there are two further regions with different scalings on the dependent variables in the interval *θ* ∈ [0, *ϵ*). Analysis within these regions is beyond the scope of this study.

Applying the above scalings to the steady-state version of Eqn. (7) and dropping the dashes, we obtain

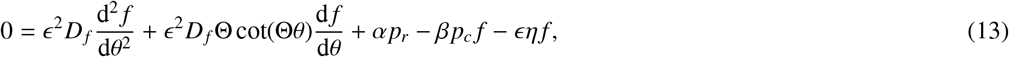

where,

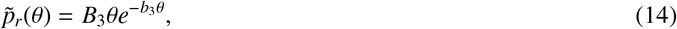

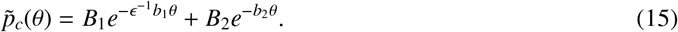

We note that cot(Θ*θ*) ≤ *O*(1) for *θ* ∈ (0.1,1] and *θ*_*r*_2__ – *θ*_*r*_1__ ≤ *O*(1). Therefore, the following analysis is valid in the region *θ* ∈ (0.1,1 – *ϵ*) (where we subtract e from the right-hand boundary to avoid the boundary layer there).

We close the system by imposing zero-flux boundary conditions as in Eqn. (11):

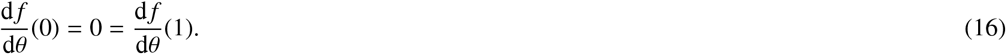

Next, we decompose the domain into left (*θ* < *θ*_*r*_1__), centre (*θ*_*r*_1__ < *θ* < *θ*_*r*_2__) and right (*θ* > *θ*_*r*_2__), outer (width *O*(1)) and inner (width *O*(*ϵ*)) regions as depicted in Fig. 3(a). Discontinuities are introduced into Eqn. (13) at the edges of the degenerate rod patch (*θ*_*r*_1__ and *θ*_*r*_2__), accounting for the left/centre/right division, while boundary layers are present either side of these discontinuities, where there is a sharp transition in *f*(*θ*), and also at the ends of the domain, to satisfy the zero-flux boundary conditions. In the following analysis we are not interested in the behaviour of the solution within the boundary layers at the ends of the domain. We label the remaining regions as follows from left to right: left-outer, left-inner, left-centre-inner, centre-outer, right-centre-inner, right-inner and right-outer (see Fig. 3(a)(ii)).

**Figure 3:**
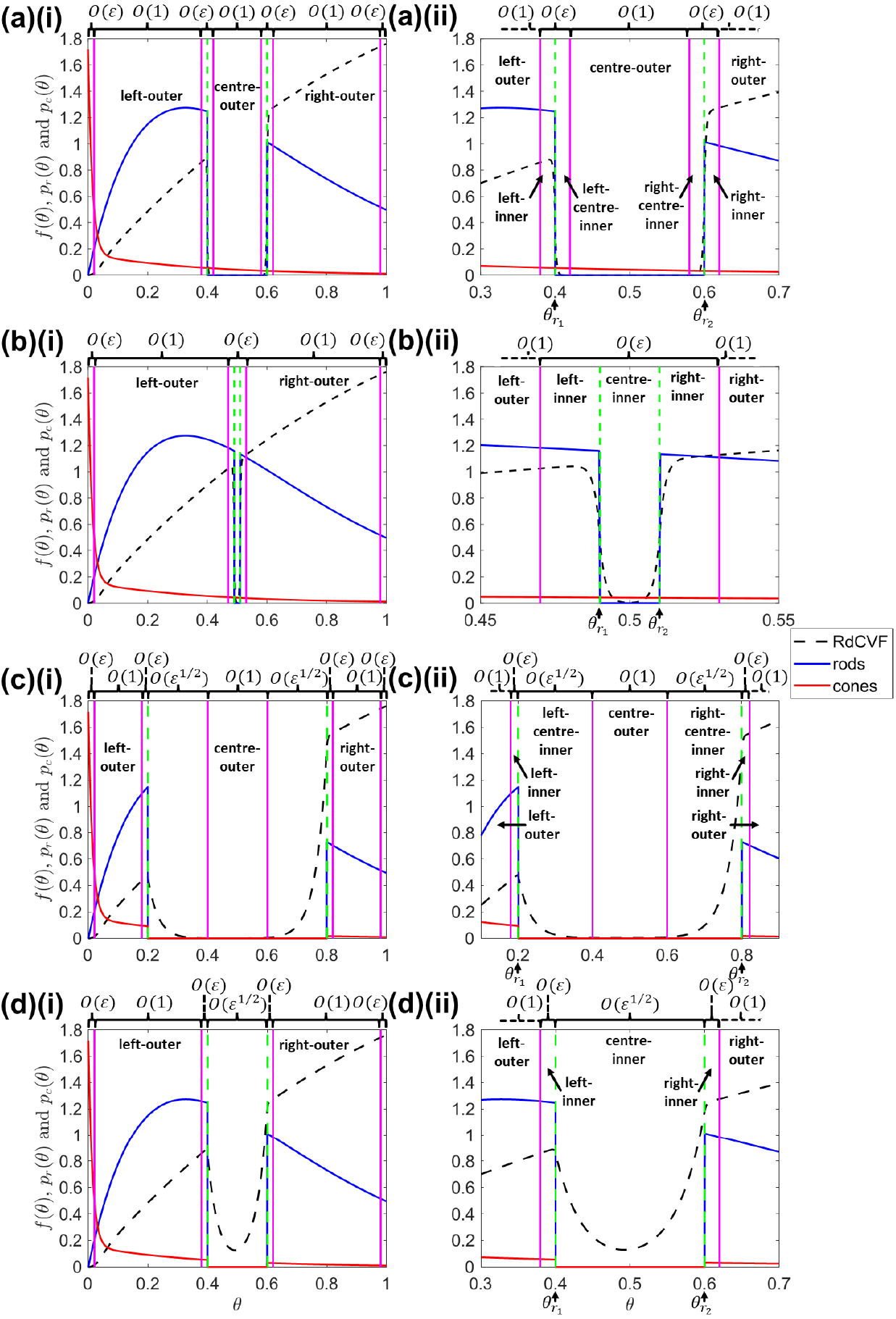
Inner and outer region locations for the asymptotic analysis. (a) wide (width *O*(1)) patch of rod loss, (b) narrow (width *O*(*ϵ*)) patch of rod loss, (c) wide (width *O*(1)) patch of rod and cone loss, (d) narrow (width *O*(*ϵ*^1/2^)) patch of rod and cone loss. The full domain is shown in the left-hand column, while the right-hand column magnifies the region from which photoreceptors have been removed. The green dashed vertical lines mark the boundaries of rod (and cone) loss, while the solid magenta vertical lines demarcate the approximate limits of the boundary layers. To obtain *f*(*θ*), Eqs. (7) and (11) were solved using the finite difference method, with 4001 mesh points, where 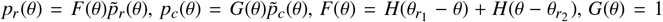 in (a) and (b), and *G*(*θ*) = *H*(*θ*_*c*_1__ – *θ*) + *H*(*θ* − *θ*_*c*_2__) in (c) and (d), and without treatment, cone regeneration or mutation-induced rod degeneration. Parameter values: *ξ* = 0; (a) (*θ*_*r*_1__, *θ*_*r*_2__) = (0.4,0.6); (b) (*θ*_*r*_1__, *θ*_*r*_2__) = (0.49,0.51); (c) (*θ*_*r*_1__, *θ*_*r*_2__) = (*θ*_*c*_1__, *θ*_*c*_2__) = (0.2,0.8); and (d) (*θ*_*r*_1__, *θ*_*r*_2__) = (*θ*_*c*_1__, *θ*_*c*_2__) = (0.4,0.6). Remaining parameter values as in Table 2.

In what follows we seek leading order (*O*(1)) solutions to *f*(*θ*) in each of the outer and inner regions. This results in a number of unknown constants of integration, whose values are determined by a combination of matching and patching. Asymptotic matching ensures that the inner limit of the outer solutions match with the outer limit of the inner solutions where they meet, while patching ensures continuity of trophic factor concentration and flux across the discontinuities at *θ*_*r*_1__ and *θ*_*r*_2__ (Bender and Orszag, 1999).

We form the regular perturbation expansions:

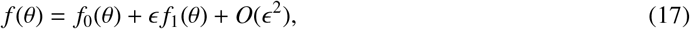

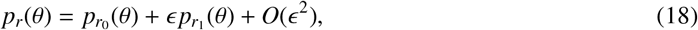

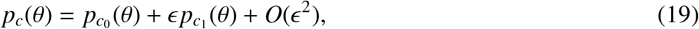

and seek *f*_0_(*θ*), where 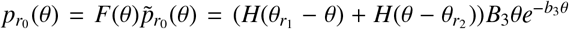 and 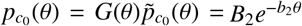.

Seeking leading order solutions to (13) in the left- and right-outer regions, we obtain,

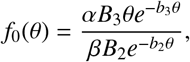

while, in the centre-outer region, we obtain,

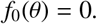

To find the leading order solution in the left-inner and left-centre-inner regions, we rescale the independent variable *θ* as 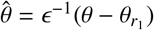, such that the first diffusion term enters the dominant balance. Thus, Eqn. (13) becomes

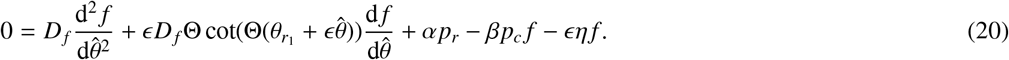

Thus, we obtain leading order solutions in the left-inner region,

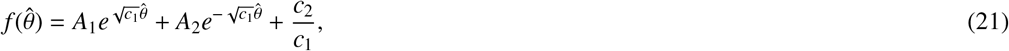

and in the left-centre-inner region,

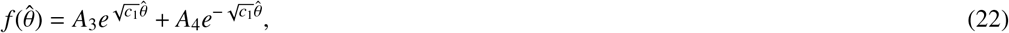

where 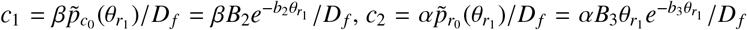, and *A*_1_–*A*_4_ are constants of integration to be determined.

To find the leading order solution in the right-centre-inner and right-inner regions, we rescale the independent variable *θ* as 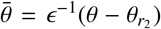, obtaining the equivalent equation to Eqn. (20), but with 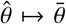 and 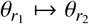.

This gives us leading order solutions in the right-centre-inner region,

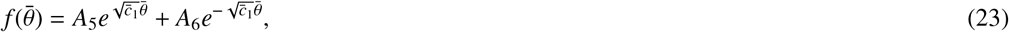

and in the right-inner region,

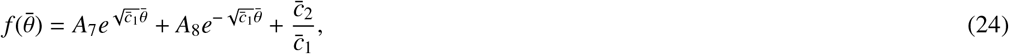

where 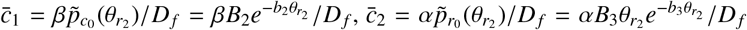, and *A*_5_–*A*_8_ are constants of integration to be determined.

To obtain values for *A*_1_–*A*_8_ we first match and then patch. We match using Van Dyke’s matching rule, which states that the *m* term inner expansion of the *n* term outer solution, should equal the *n* term outer expansion of the *m* term inner solution. In this case, since we are dealing with leading order solutions only, *m* = *n* = 1. Matching the left-outer and left-inner solutions, we form the one term inner expansion of the one term outer solution,

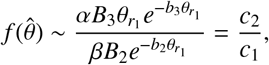

and the one term outer expansion of one term inner solution,

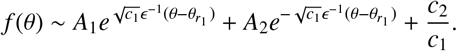

Since *θ* – *θ*_*r*_1__ > 0 in the left-hand regions, the first term is exponentially small and so can be neglected, while the second term is exponentially large and so we must have *A*_2_ = 0. Matching the left-centre-inner and centre-outer solutions, the centre-outer and right-centre-inner solutions, and the right-inner and right-outer solutions in a similar fashion, we obtain *A*_3_ = 0, *A*_6_ = 0, and *A*_7_ = 0 respectively.

Patching across *θ* = *θ*_*r*_1__ and *θ* = *θ*_*r*_2__, we obtain expression for *A*_1_, *A*_4_, *A*_5_ and *A*_8_ in terms of known parameters. Substituting for *A*_1_–*A*_8_ into Eqs. (21)–(24), we obtain the full set of leading order outer and inner solutions as follows —

Left- and Right-Outer:

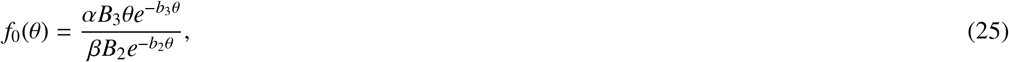

Left-Inner:

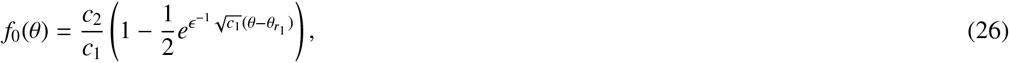

Left-Centre-Inner:

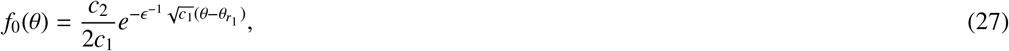

Centre-Outer:

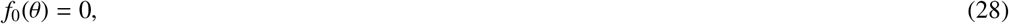

Right-Centre-Inner:

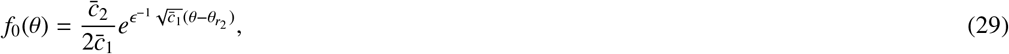

Right-Inner:

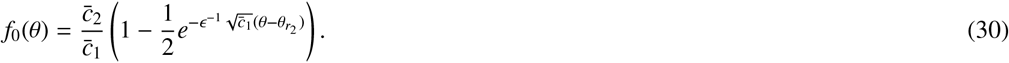

Next we calculate leading order composite solutions, providing uniformly valid expansions across the left-outer and left-inner regions (left-composite); across the left-centre-inner, centre-outer and right-centre-inner regions (centre-composite), and across the right-inner and right-outer regions (right-composite). Composite solutions are calculated by adding together the relevant outer and inner solutions and subtracting one copy of the common terms to avoid double counting (see Bender and Orszag, 1999, for further details). We obtain the following composite solutions —

Left-Composite:

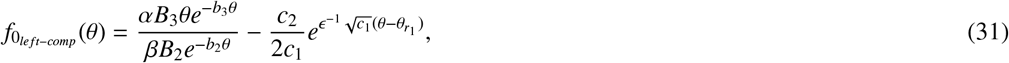

Centre-Composite:

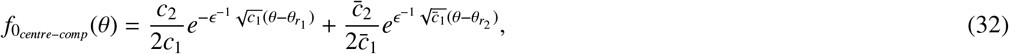

Right-Composite:

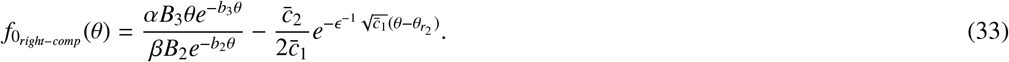

Lastly, we derive expressions for the cone degenerate patch boundaries. As discussed above, we assume that the boundaries of a cone degenerate patch, *θ*_*crit*_1__ (left-hand boundary) and *θ*_*crit*_2__ (right-hand boundary), occur where *f*(*θ*) = *f_crit_*. This cannot occur in the outer regions, since we have chosen *f_crit_* such that *f*(*θ*) > *f_crit_* in the healthy retina (left- and right-outer regions), while *f*_0_(*θ*) = 0 < *f_crit_* in the centre-outer region. Therefore, we must have *f*(*θ*) = *f_crit_* somewhere in the left-inner or left-centre-inner region (*θ*_*crit*_1__), and somewhere in the right-centre-inner or right-inner region (*θ*_*crit*_2__). Setting *f*(*θ*) = *f_crit_* in Eqs. (26), (27), (29) and (30), and rearranging, we obtain —

Minimal Cone Degenerate Patch Left Boundary:

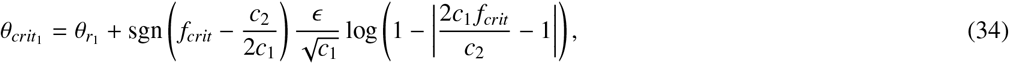

Maximal Cone Degenerate Patch Right Boundary:

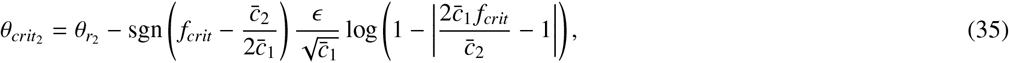

where sgn(·) is the signum (or sign) function, defined such that,

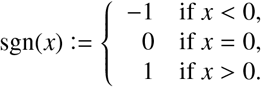

We have combined the expressions for *θ*_*crit*_1__ resulting from Eqs. (26) and (27), to give Eqn. (34), and similarly for *θ*_*crit*_2__ with Eqs. (29) and (30) to give Eqn. (35). The position of the cone degenerate patch boundaries relative to the rod degenerate patch boundaries, and hence the question of whether the cone degenerate patch lies within or exceeds the rod degenerate patch, depends upon the value of *f_crit_*, and is given by the following inequalities —

- when *f_crit_* ≥ *c*_2_/(2*c*_1_), *θ*_*crit*_1__ ≤ *θ*_*r*_1__;
- when *f_crit_* ≥ *c*_2_/(2*c*_1_), *θ*_*crit*_1__ > *θ*_*r*_1__;
- when 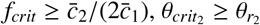
- when 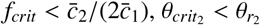.

Thus, as would be expected intuitively, cone degenerate patches are wider for larger values of *f_crit_*.

For asymptotic analysis Cases 2-8, see Appendix A.

### 3.2. Numerical and analytical solutions

In this section we compare numerical solutions and analytical approximations to the steady-state problem in which rods (or rods and cones) have been lost from the interval *θ* ∈ (*θ*_*r*_1__, *θ*_*r*_2__), with and without treatment. Analytical solutions provide deeper insight into the dominant mechanisms determining the behaviour of our models and are cheap to compute, while numerical solutions serve to confirm their accuracy and are more computationally expensive. To obtain numerical solutions we solve Eqs. (7) and (11) for *f*(*θ*) at steady-state using the Matlab routine fsolve, which employs a Trust–Region–Dogleg algorithm. Analytical approximations are derived in Section 3.1 and Appendix A.

We consider the 8 cases described in Section 3.1, dividing each of these cases into the two subcases:

1. Low (homogeneous) RdCVF threshold concentration *f_crit_* = 3 × 10^−5^: chosen such that *f_crit_* < *f*(*θ*) for all *θ* ∈ [0,1] in the healthy retina (where 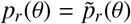 and 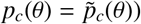;
2. High (heterogeneous) RdCVF threshold concentration *f_crit_* = *f_crit_*(*θ*) = *f*_*crit*_1__ *H*(0.13 − *θ*) + *f*_*crit*_2__ (1 – *H*(0.13 − *θ*)): where *f*_*crit*_1__ = 3 × 10^−5^, *f*_*crit*_2__ = 0.3 and the switch point, *θ* = 0.13, are chosen such that *f_crit_*(*θ*) < *f*(*θ*) for all *θ* ∈ [0,1] in the healthy retina.

The value of *f_crit_* has not yet been measured; therefore, we consider the two most likely scenarios. In the first (low *f_crit_*) scenario, the threshold is spatially uniform and chosen to be low enough such that cones will not degenerate in the healthy retina. In the second (high *f_crit_*) scenario, we assume that the cone-rich fovea is afforded special protection against low RdCVF levels, such that *f_crit_* is lower in that region (*θ* ≤ 0.13) than in the rest of the retina, where rods, and hence RdCVF, are more plentiful and cone density, and hence the rate of RdCVF uptake, is low. For notational simplicity, we shall refer to the high *f_crit_* scenario simply as *f_crit_* = 0.3 in what follows. Note that throughout this section, we consider only the region *θ* ∈ (0.13,1] in the high *f_crit_* scenario.

#### 3.2.1. No treatment

We begin by considering the untreated case, in which *ξ* = 0. Here we are interested in predicting the interval over which cones will degenerate following the removal of a patch of rods from the interval *θ* ∈ (*θ*_*r*_1__, *θ*_*r*_2__), and the conditions under which cone degeneration will expand beyond the rod degenerate patch. The boundaries of the resultant patch of cone loss are calculated as the positions at which *f*(*θ*) = *f_crit_* (between which *f* (*θ*) < *f_crit_*) local to the patch of rod loss. In the full dynamic problem (see Section 4.1), *f*(*θ*) rises as cones degenerate, shrinking the interval over which *f*(*θ*) < *f_crit_*. Here, we derive upper and lower bounds on the cone degenerate patch width. The upper bound is found in the case where only rods are removed, such that cones continue to consume RdCVF within the degenerate patch, expanding the interval over which *f*(*θ*) < *f_crit_*. The lower bound is found in the case where both rods and cones are removed from *θ* ∈ (*θ*_*r*_1__, *θ*_*r*_2__) (where *θ*_*c*_1__ = *θ*_*r*_1__ and *θ*_*c*_2__ = *θ*_*r*_2__), such that cones do not consume RdCVF within the degenerate patch, reducing the interval over which *f* (*θ*) < *f_crit_*. We note that, in calculating the lower bound this way, we assume that cones do not degenerate beyond the rod degenerate patch. Where cone loss does exceed the rod patch, this method acts as a confirmation that this will occur, rather than as a lower bound.

Fig. 4 shows analytical solutions for the distance between the left-hand rod and cone degenerate patch boundaries, *θ*_*r*_1__ – *θ*_*crit*_1__, as a function of *θ*_*r*_1__, and similarly for the distance between the right-hand rod and cone degenerate patch boundaries, *θ*_*crit*_2__ – *θ*_*r*_2__, as a function of *θ*_*r*_2__. We consider the rod loss only ((a) and (b)), and rod and cone loss cases ((c) and (d)), corresponding to the upper and lower bounds on cone degenerate patch width, respectively. For each of these cases, we consider the subcases *f_crit_* = 0.3 ((a) and (c)) and *f_crit_* = 3 × 10^−5^ ((b) and (d)).

**Figure 4:**
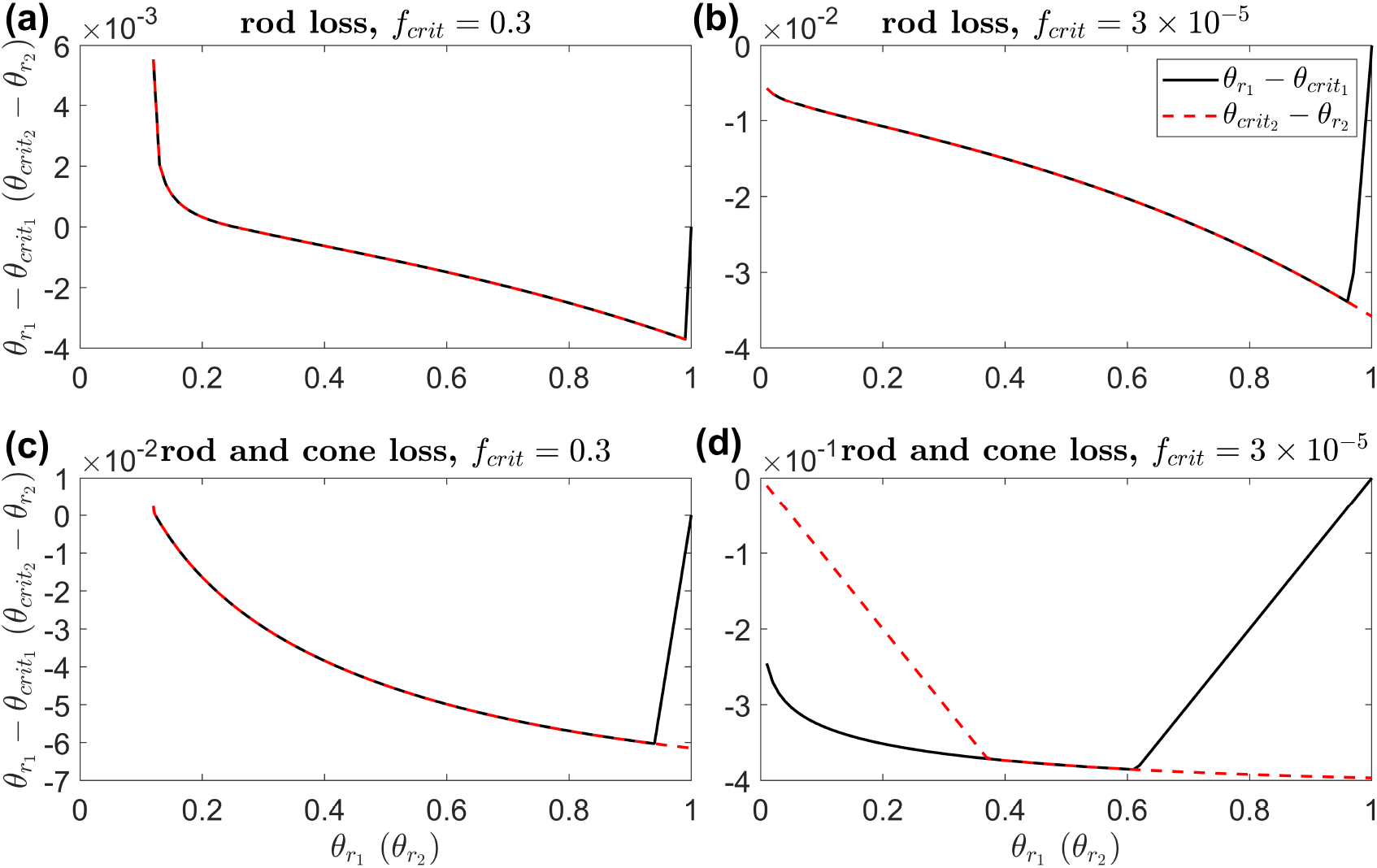
Distance between rod and minimum/maximum cone degenerate patch boundaries — wide rod degenerate patch case. Each panel shows the variation in the distances between the left-hand rod and minimum cone degenerate patch boundaries, *θ*_*r*_1__ – *θ*_*crit*_1__, and between the right-hand rod and maximum cone degenerate patch boundaries, *θ*_*crit*_2__ – *θ*_*r*_2__, with *θ*_*r*_1__ and *θ*_*r*_2__ respectively. (a) rod loss only with *f_crit_* = 0.3, (b) rod loss only with *f_crit_* = 3 × 10^−5^, (c) rod and cone loss with *f_crit_* = 0.3, (d) rod and cone loss with *f_crit_* = 3 × 10^−5^. Note the different scales on the y-axes. The maximum spatial extent of cone loss remains within the boundaries of rod loss in (b) and (d), but may exceed it close to the fovea (centred at *θ* = 0) in (a) and (c). Curves are plotted using Eqs. (34) and (35) in (a) and (b), and using Eqs. (A.21)–(A.24) in (c) and (d). The parameter β = 10^−2^. Remaining parameter values as in Table 2.

The equations plotted in Fig. 4 (*θ*_*r*_1__ – *θ*_*crit*_1__ and *θ*_*crit*_2__ – *θ*_*r*_2__, where *θ*_*crit*_1__ and *θ*_*crit*_2__ are as given in Eqs. (34) and (35) in (a) and (b), and Eqs. (A.21)–(A.24) in (c) and (d)) are all monotone decreasing functions of *θ*_*r*_1__ and *θ*_*r*_2__. However, *θ*_*crit*_1__ and *θ*_*crit*_2__ cannot in practice go outside of the retina and hence cannot exceed the interval *θ* ∈ [0,1]. Therefore, we set *θ*_*crit*_1__ = 1 where *θ*_*crit*_1__ > 1 and *θ*_*crit*_2__ = 0 where *θ*_*crit*_2__ < 1. This results in upward jumps on the right-hand side of the domain in θ*r*_1_ – θ*crit*_1_ in (a)–(d) and an upward jump on the left-hand side of the domain in *θ*_*crit*_2__ – *θ*_*r*_2__ in (d). Apart from these jumps, the curves for *θ*_*r*_1__ – *θ*_*crit*_1__ and *θ*_*crit*_2__ – *θ*_*r*_2__ are identical. Note that, for the wide patch case, the predicted position of one cone degenerate patch boundary does not depend upon the other cone degenerate patch boundary (unlike in the narrow patch case), since the boundary layers surrounding the rod degenerate patch boundaries are separated by a central outer region, such that they do not ‘interact’. The maximum spatial extent of cone loss remains within the boundaries of rod loss in (b) and (d), but may exceed it close to the fovea (centred at *θ* = 0) in (a) and (c).

Fig. 5 shows numerical and analytical bounds on cone degenerate patch size for the wide rod degenerate patch case. The cone degenerate patch width, *θ*_*crit*_2__ – *θ*_*crit*_1__, increases monotonically with increasing rod degenerate patch width, *θ*_*r*_2__ – *θ*_*r*_1__, and with decreasing rod degenerate patch centre position, (*θ*_*r*_1__ + *θ*_*r*_2__)/2. For narrower rod loss patches in (c), and especially (d), the predicted lower bound on cone degenerate patch width is zero. The difference between rod and cone degenerate patch widths, (*θ*_*crit*_2__ – *θ*_*crit*_1__) – (*θ*_*r*_2__ – *θ*_*r*_1__), is a monotone (not strictly) decreasing function of rod degenerate patch centre position. The cone degenerate patch width is less than the rod degenerate patch width except in (a) (rod loss only with *f_crit_* = 0.3), where it may exceed rod degenerate patch width close to the fovea, doing so for a wider range of patch positions for narrower rod loss patches. The predicted cone degenerate patch width is lower for the rod and cone loss case ((a) and (b), lower bound) than for the rod loss only case ((c) and (d), upper bound) as would be expected. Numerical and analytical results match closely, except for the width 0.1 rod and cone degenerate patch case in (c), where the wide patch assumption breaks down, since 0.1 ~ *O*(*ϵ*^1/2^).

**Figure 5:**
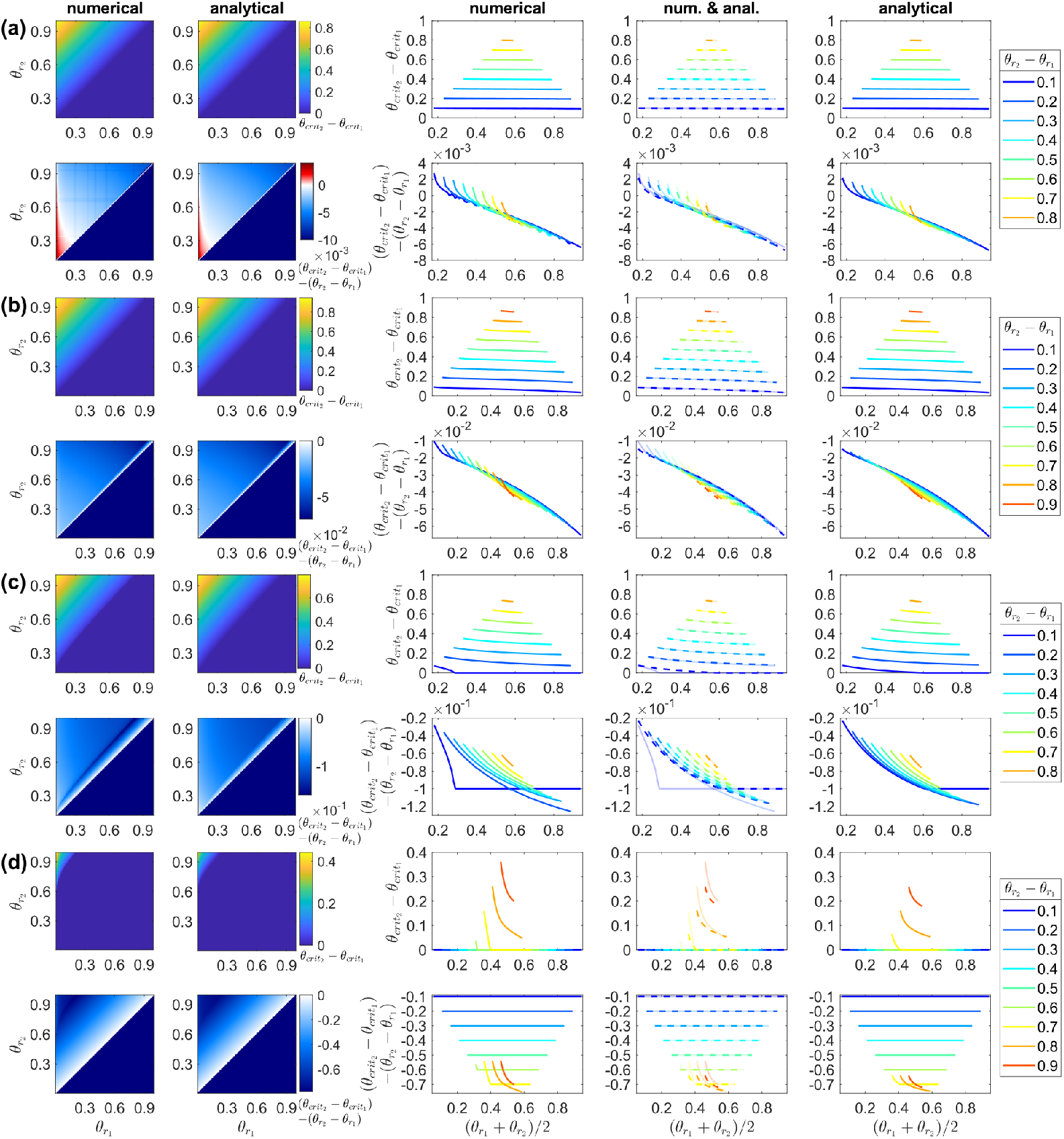
Numerical and analytical bounds on cone degenerate patch size using the steady-state model — wide rod degenerate patch case. (a)–(d) first row: maximum/minimum cone degenerate patch width, *θ*_*crit*_2__ – *θ*_*crit*_1__; second row: difference between rod and maximum/minimum cone degenerate patch widths, (*θ*_*crit*_2__ – *θ*_*crit*_1__) – (*θ*_*r*_2__ – *θ*_*r*_1__); columns 1 and 2: variation of patch widths and width differences in (*θ*_*r*_1__, *θ*_*r*_2__) parameter space; columns 3–5: variation of patch widths and width differences with rod degenerate patch centre position, (*θ*_*r*_1__ + *θ*_*r*_2__)/2, each curve representing a constant rod degenerate patch width, *θ*_*r*_2__ – *θ*_*r*_1__; columns 1 and 3: numerical solutions; columns 2 and 5: analytical approximations; column 4: numerical (solid curves) and analytical (dashed curves) solutions compared. (a) rod loss only with *f_crit_* = 0.3, (b) rod loss only with *f_crit_* = 3 × 10^−5^, (c) rod and cone loss with *f_crit_* = 0.3, (d) rod and cone loss with *f_crit_* = 3 × 10^−5^. (a) and (b) upper bounds on cone degenerate patch widths; (c) and (d) lower bounds on cone degenerate patch widths. Cone degenerate patch width increases monotonically with increasing rod degenerate patch width and with decreasing rod degenerate patch centre position. Cone degenerate patch width is less than rod degenerate patch width except in (a), where it may exceed rod degenerate patch width close to the fovea (centred at *θ* = 0). Numerical solutions were obtained by calculating the eccentricities at which *f*(*θ*) = *f_crit_*, solving Eqs. (7) and (11) for *f*(*θ*) at steady-state using the finite difference method, with 4001 mesh points, where 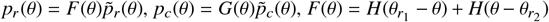, *G*(*θ*) = 1 in (a) and (b), and *G*(*θ*) = *H*(*θ*_*c*_1__ – *θ*) + *H*(*θ* – *θ*_*c*_2__), with *θ*_*c*_1__ = *θ*_*r*_1__ and *θ*_*c*_2__ = *θ*_*r*_2__, in (c) and (d), and without treatment or cone regeneration. Analytical solutions are plotted using Eqs. (34) and (35) in (a) and (b), and using Eqs. (A.21)–(A.24) in (c) and (d). Parameter values: *ξ* = 0 and *ϵ* = 10^−2^. Remaining parameter values as in Table 2.

Fig. 6 shows numerical and analytical bounds on cone degenerate patch size for the narrow rod degenerate patch case. We plot only the *f_crit_* = 0.3 subcase here since, for the *f_crit_* = 3 × 10^−5^ subcase, the predicted cone loss patch width is always zero. As in Fig. 5, cone degenerate patch width increases monotonically with increasing rod degenerate patch width and with decreasing rod degenerate patch centre position, while the difference between rod and cone degenerate patch widths is a monotone (not strictly) decreasing function of rod degenerate patch centre position. The cone degenerate patch width may exceed the rod degenerate patch width close to the fovea in (a). Numerical and analytical solutions agree well in (a) and on the right-hand side in (b); however, the scaling assumptions break down close to the fovea, such that the analytical solution predicts cone loss on left-hand side in (b), where the numerical solution remains zero.

**Figure 6:**
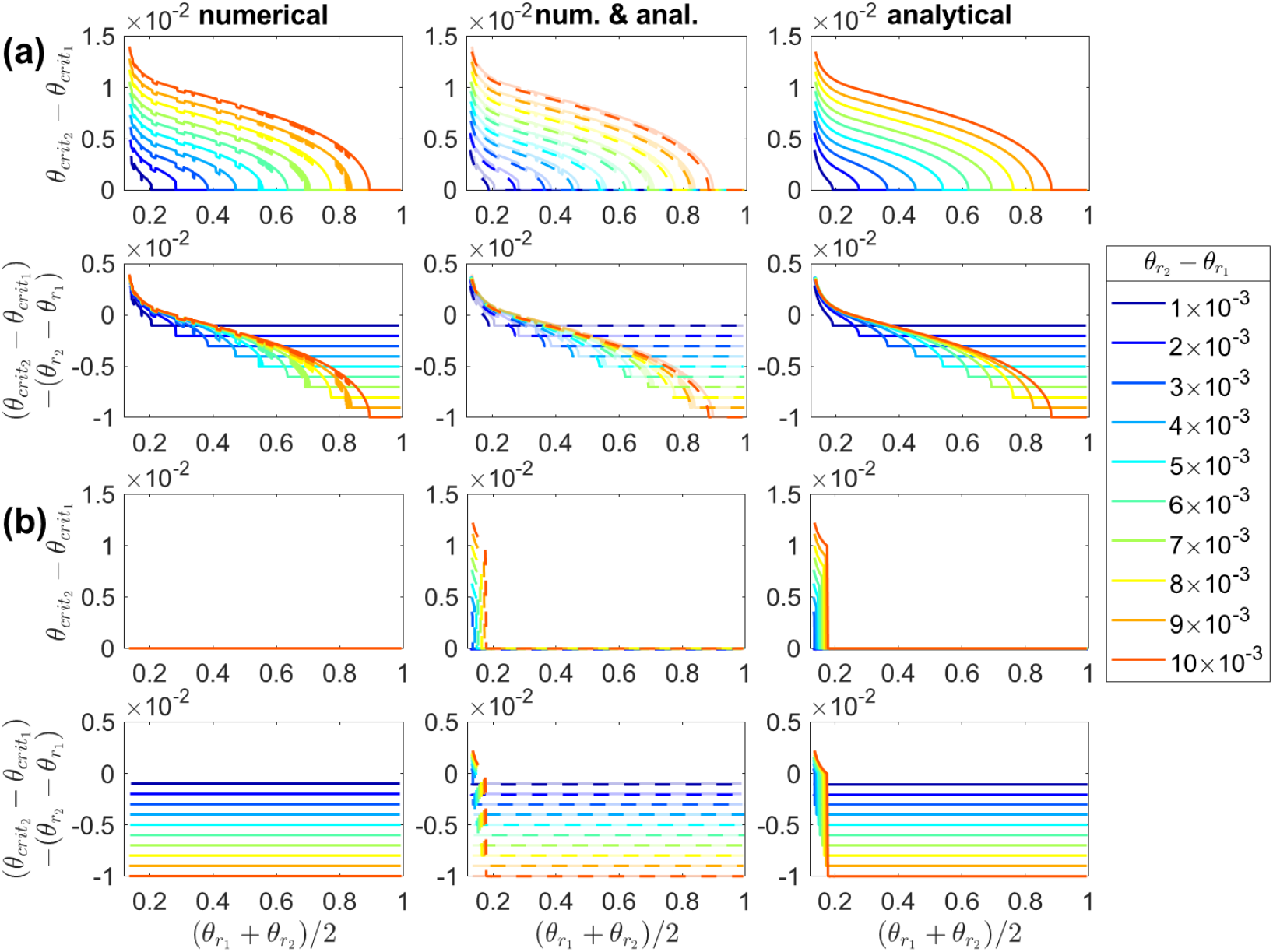
Numerical and analytical bounds on cone degenerate patch size using the steady-state model — narrow rod degenerate patch case. Graphs show variation of maximum/minimum cone degenerate patch widths, and differences between rod and maximum/minimum cone degenerate patch widths with rod degenerate patch centre position, (*θ*_*r*_1__ + *θ*_*r*_2__)/2, each curve representing a constant rod degenerate patch width, *θ*_*r*_2__ – *θ*_*r*_1__. (a) and (b) first row: maximum/minimum cone degenerate patch width, *θ*_*crit*_2__ – *θ*_*crit*_1__ second row: difference between rod and maximum/minimum cone degenerate patch widths, (*θ*_*crit*_2__ – *θ*_*crit*_1__) – (*θ*_*r*_2__ – *θ*_*r*_1__); column 1: numerical solutions; column 3: analytical approximations; column 2: numerical (solid curves) and analytical (dashed curves) solutions compared. (a) rod loss only: upper bounds on cone degenerate patch widths; (b) rod and cone loss: lower bounds on cone degenerate patch widths. Cone degenerate patch width increases monotonically with increasing rod degenerate patch width and with decreasing rod degenerate patch centre position. Cone degenerate patch width may exceed rod degenerate patch width close to the fovea (centred at *θ* = 0). Numerical solutions were obtained by calculating the eccentricities at which f(*θ*) = *f_crit_*, solving Eqs. (7) and (11) for *f*(*θ*) at steady-state using the finite difference method, with 4001 mesh points, where 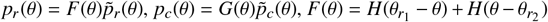, *G*(*θ*) = 1 in (a), and *G*(*θ*) = *H*(*θ*_*c*_1__ – *θ*) + *H*(*θ* – *θ*_*c*_2__), with *θ*_*c*_1__ = *θ*_*r*_1__ and *θ*_*c*_2__ = *θ*_*r*_2__, in (b), and without treatment or cone regeneration. Analytical solutions are plotted using Eqs. (A.7)–(A.10) in (a), and using Eqs. (A.31)–(A.34) in (b). Parameter values: *f_crit_* = 0.3, *ξ* = 0 and *ϵ* = 10^−2^. Remaining parameter values as in Table 2.

#### 3.2.2. Treatment

In this section we compare numerical and analytical estimates for the critical treatment rate, *ξ_crit_*. This is the minimum RdCVF treatment rate required to prevent cone degeneration in a given scenario. We consider just the rod loss only case, since this provides an upper bound on *ξ_crit_*, which is of greater clinical relevance, given that this would theoretically guarantee the avoidance of cone degeneration under the trophic factor hypothesis. We also consider analytical estimates for the eccentricity of the minimum trophic factor concentration local to the rod degenerate patch, *θ_crit_*, when *ξ* = *ξ_crit_*. This is the point at which cone loss will initiate if drops below *ξ_crit_*. Numerical solutions are obtained using the Matlab routine fminsearch (which uses a simplex search method) to find the value of at which the minimum value of *f*(*θ*) local to the rod degenerate patch is equal to *f_crit_*. This involves solving Eqs. (7) and (11) for *f*(*θ*) at steady-state using fsolve at each iteration. Analytical solutions are obtained by employing the Matlab routine fsolve to solve the implicit equations (A.46) and (A.47) for a wide patch of rod loss with global treatment, (A.59) and (A.60) for a wide patch of rod loss with local treatment, (A.71) and (A.72) for a narrow patch of rod loss with global treatment, and (A.83) and (A.84) for a narrow patch of rod loss with local treatment.

Fig. 7 shows numerical and analytical estimates of the critical treatment rate for the wide rod degenerate patch case. The critical treatment rate increases monotonically with decreasing rod degenerate patch centre position and left boundary position in all cases. The critical treatment rate depends almost exclusively upon the rod degenerate patch left boundary position, rather than the rod degenerate patch width or right boundary position. The critical treatment rate is *O*(10^4^) for *f_crit_* = 0.3 ((a) and (b)) and *O*(1) for *f_crit_* = 3 × 10^−5^ ((c) and (d)), a lower rate being sufficient to raise the trophic factor concentration above the lower critical threshold. Numerical and analytical solutions agree well.

**Figure 7:**
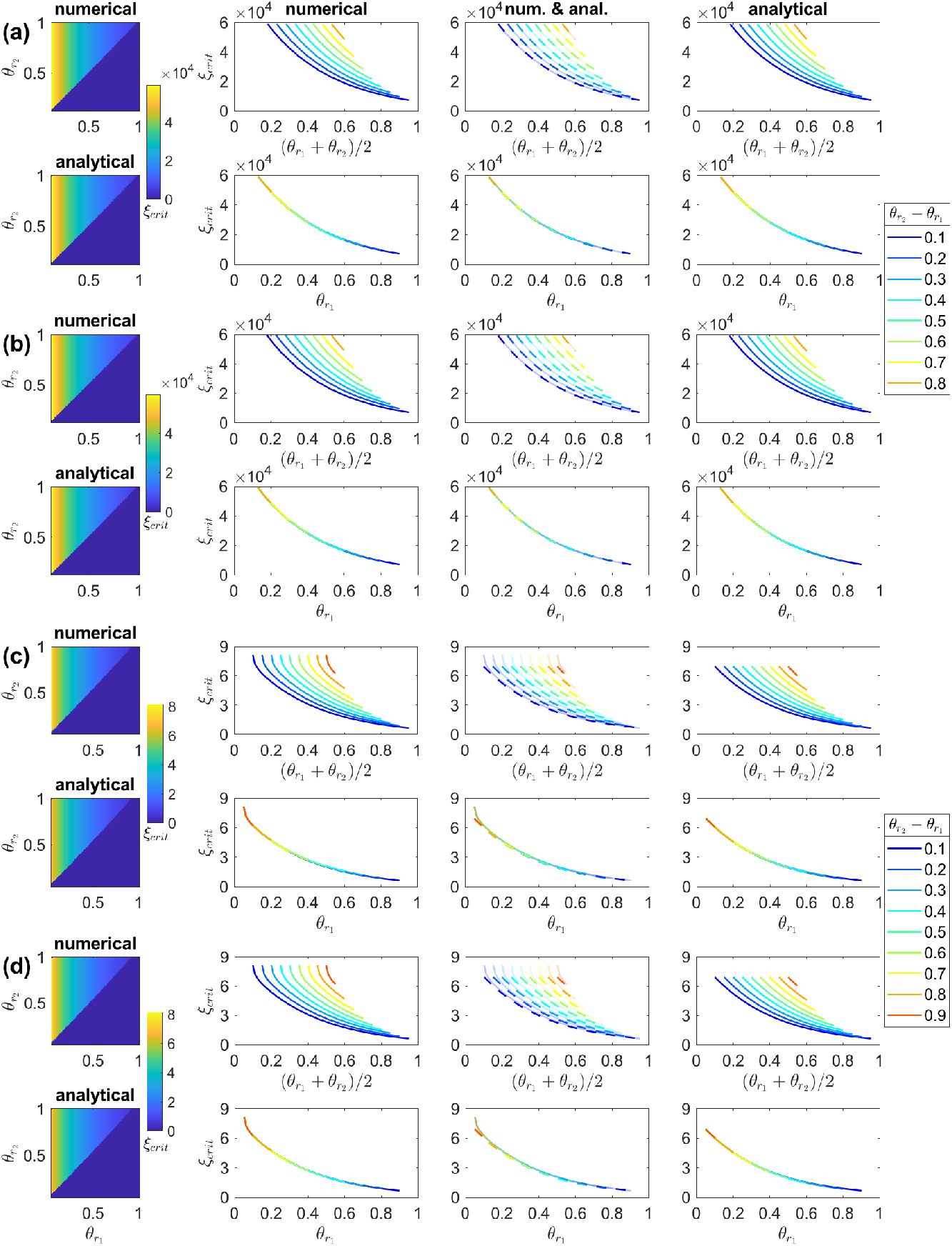
Numerical and analytical estimates of the critical treatment rate using the steady-state model — wide rod degenerate patch case. Rod loss only. All plots show the critical treatment rate, *ξ_crit_*. (a) and (c) global treatment, (b) and (d) local treatment; (a) and (b) *f_crit_* = 0.3; (c) and (d) *f_crit_* = 3 × 10^−5^. (a)–(d) column 1: numerical (top) and analytical (bottom) estimates of the variation in the critical treatment rate over (*θ*_*r*_1__, *θ*_*r*_2__) parameter space; columns 2–4: variation in the critical treatment rate with rod degenerate patch centre position, (*θ*_*r*_1__ + *θ*_*r*_2__)/2 (top), and with rod degenerate patch left boundary position, *θ*_*r*_1__ (bottom), each curve representing a constant rod degenerate patch width, *θ*_*r*_2__ – *θ*_*r*_1__; column 2: numerical solutions; column 4: analytical approximations; column 3: numerical (solid curves) and analytical (dashed curves) solutions compared. The critical treatment rate increases monotonically with decreasing rod degenerate patch centre/left boundary position in all cases. The critical treatment rate depends almost entirely upon the rod degenerate patch left boundary position, rather than the rod degenerate patch width/right boundary position. Numerical solutions were obtained by using the Matlab routine fminsearch to calculate the critical treatment rate at which min(*f*(*θ*)) = *f_crit_* within *θ* ∈ [*θ*_*r*_1__, *θ*_*r*_2__], solving Eqs. (7) and (11) for *f*(*θ*) at steady-state at each iteration using the finite difference method, with 401 mesh points, where 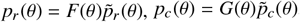, *F*(*θ*) = *H*(*θ*_*r*_1__ – *θ*) + *H*(*θ* – *θ*_*r*_2__), *G*(*θ*) = 1, *T*(*θ*) = 1 for (a) and (c), and *T*(*θ*) = 1 – *F*(*θ*) for (b) and (d). Analytical solutions are obtained by implicitly solving Eqs. (A.46) and (A.47) in (a) and (c), and Eqs. (A.59) and (A.60) in (b) and (d). The parameter *ϵ* = 10^−2^. Remaining parameter values as in Table 2.

Fig. 8 shows numerical and analytical estimates of the critical treatment rate for the narrow rod degenerate patch case. We consider only the *f_crit_* = 0.3 subcase here, since there is no cone loss for *f_crit_* = 3 × 10^−5^ and hence no treatment is required in this subcase. As with the wide rod degenerate patch case, the critical treatment rate increases monotonically with decreasing rod degenerate patch centre position and left boundary position in all cases. Unlike in the wide rod degenerate patch case, the critical treatment rate depends upon both the rod degenerate patch left and right boundary positions, and hence upon the patch width (compare Fig. 7), increasing monotonically with increasing rod degenerate patch width. Numerical and analytical solutions match closely.

**Figure 8:**
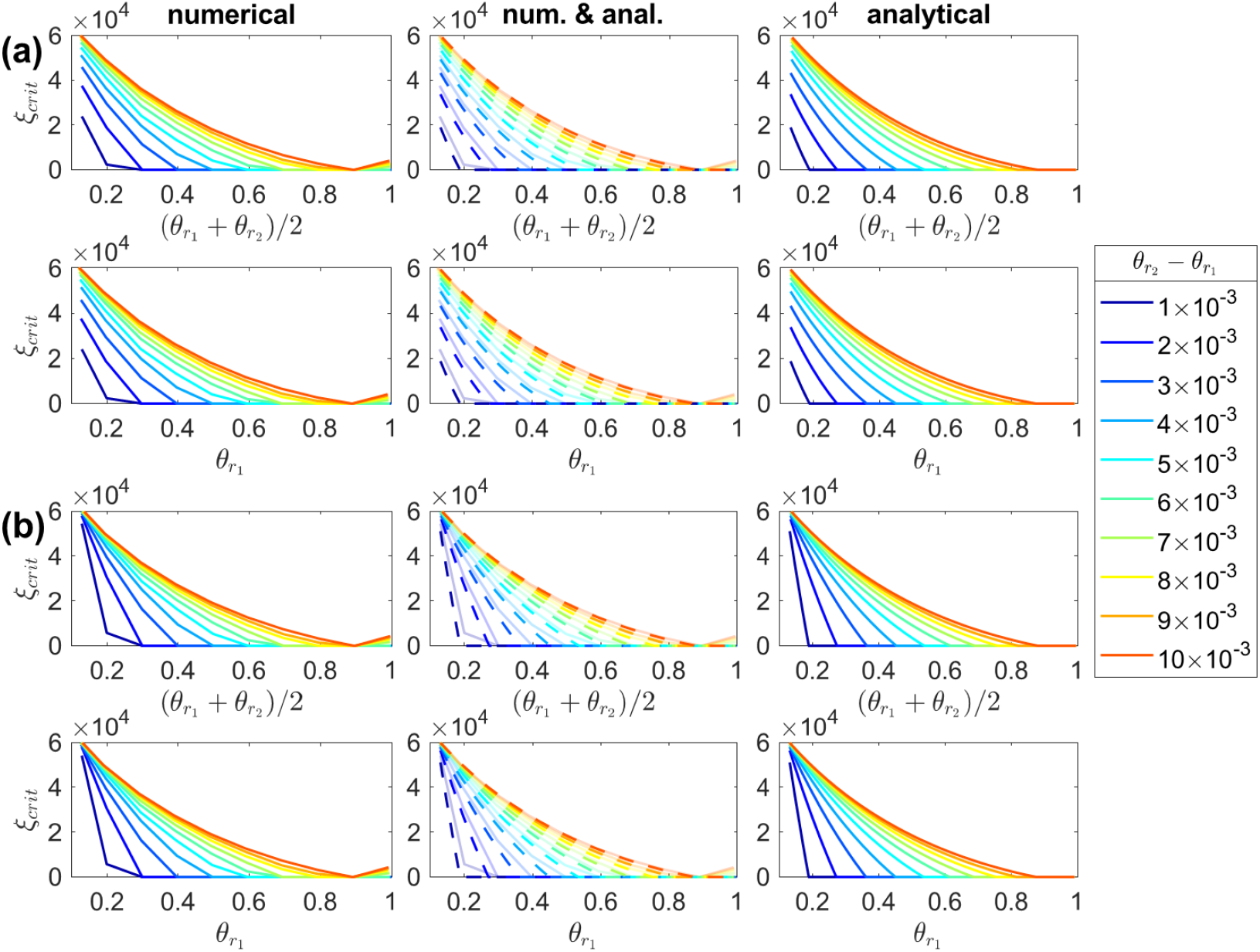
Numerical and analytical estimates of the critical treatment rate using the steady-state model — narrow rod degenerate patch case. Rod loss only. All plots show the critical treatment rate, *ξ_crit_*. (a) global treatment, (b) local treatment. (a) and (b) variation in the critical treatment rate with rod degenerate patch centre position, (*θ*_*r*_1__ + *θ*_*r*_2__)/2 (top), and with rod degenerate patch left boundary position, *θ*_*r*_1__ (bottom), each curve representing a constant rod degenerate patch width, *θ*_*r*_2__ – *θ*_*r*_1__; column 1: numerical solutions; column 3: analytical approximations; column 2: numerical (solid curves) and analytical (dashed curves) solutions compared. The critical treatment rate depends upon both the rod degenerate patch left and right boundary positions, and hence the patch width (compare Fig. 7). The critical treatment rate increases monotonically with increasing rod degenerate patch width and with decreasing patch centre/left boundary position in all cases. Numerical solutions were obtained by using the Matlab routine fminsearch to calculate the critical treatment rate at which min(*f*(*θ*)) = *f_crit_* within *θ* ∈ [*θ*_*r*_1__, *θ*_*r*_2__], solving Eqs. (7) and (11) for *f*(*θ*) at steady-state at each iteration using the finite difference method, with 1001 mesh points, where 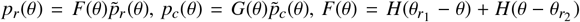, *G*(*θ*) = 1, *T*(*θ*) = 1 for (a) and *T*(*θ*) = 1 – *F*(*θ*) for (b). Analytical solutions are obtained by implicitly solving Eqs. (A.71) and (A.72) in (a) and Eqs. (A.83) and (A.84) in (b). Parameter values: *f_crit_* = 0.3 and β = 10^−2^. Remaining parameter values as in Table 2.

Fig. 9 compares analytical estimates of the critical treatment rate for the global and local treatment cases. The critical treatment rate is almost identical for global and local treatment in the wide patch case, for both *f_crit_* = 0.3 (a) and *f_crit_* = 3 × 10^−5^ (b), while the critical treatment rate is higher for local than for global treatment for the narrowest patches close to the fovea (centred at *θ* = 0) in the narrow patch case (with *f_crit_* = 0.3) (c).

**Figure 9:**
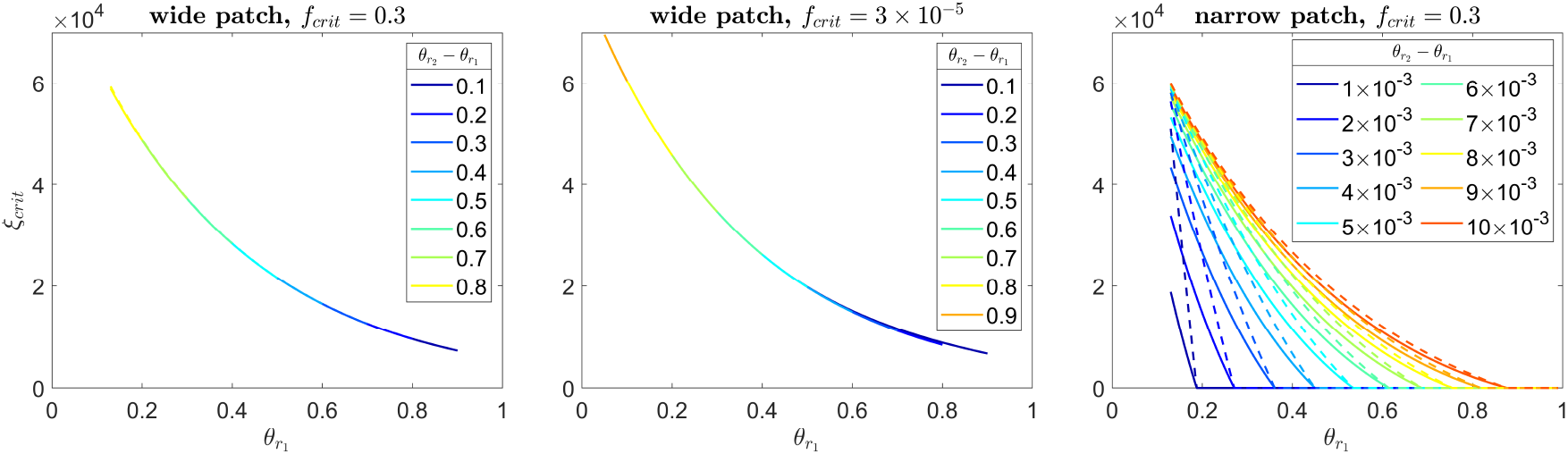
Comparison of critical treatment rates under global and local treatment — wide and narrow rod degenerate patch cases. All plots show analytical estimates of the variation in critical treatment rate, *ξ_crit_*, with rod degenerate patch left boundary position, *θ*_*r*_1__, using the steady-state model. Each curve represents a constant rod degenerate patch width, *θ*_*r*_2__ – *θ*_*r*_1__. Solid curves: global treatment; dashed curves: local treatment. (a) wide rod degenerate patch with *f_crit_* = 0.3, (b) wide patch with *f_crit_* = 3 × 10^−5^, (c) narrow patch with *f_crit_* = 0.3. The critical treatment rate is almost identical for global and local treatment in the wide patch case, while the critical treatment rate is higher for local than for global treatment for the narrowest patches close to the fovea (centred at *θ* = 0) in the narrow patch case. Analytical solutions and parameter values are identical to those for the corresponding cases in Figs. 7 and 8.

Fig. 10 shows analytical estimates of the normalised eccentricity of the minimum trophic factor concentration, 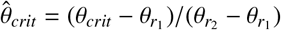, for both the wide and narrow rod degenerate patch cases. The variable 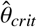 increases monotonically with decreasing rod degenerate patch width, and with increasing rod degenerate patch centre position and left boundary position in all cases. Further, 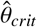 remains close to 0 for patches of width *θ*_*r*_2__ – *θ*_*r*_1__ ≥ 0.2 ((a)–(d)) and close to 0.5 for narrow patches ((e) and (f)).

**Figure 10:**
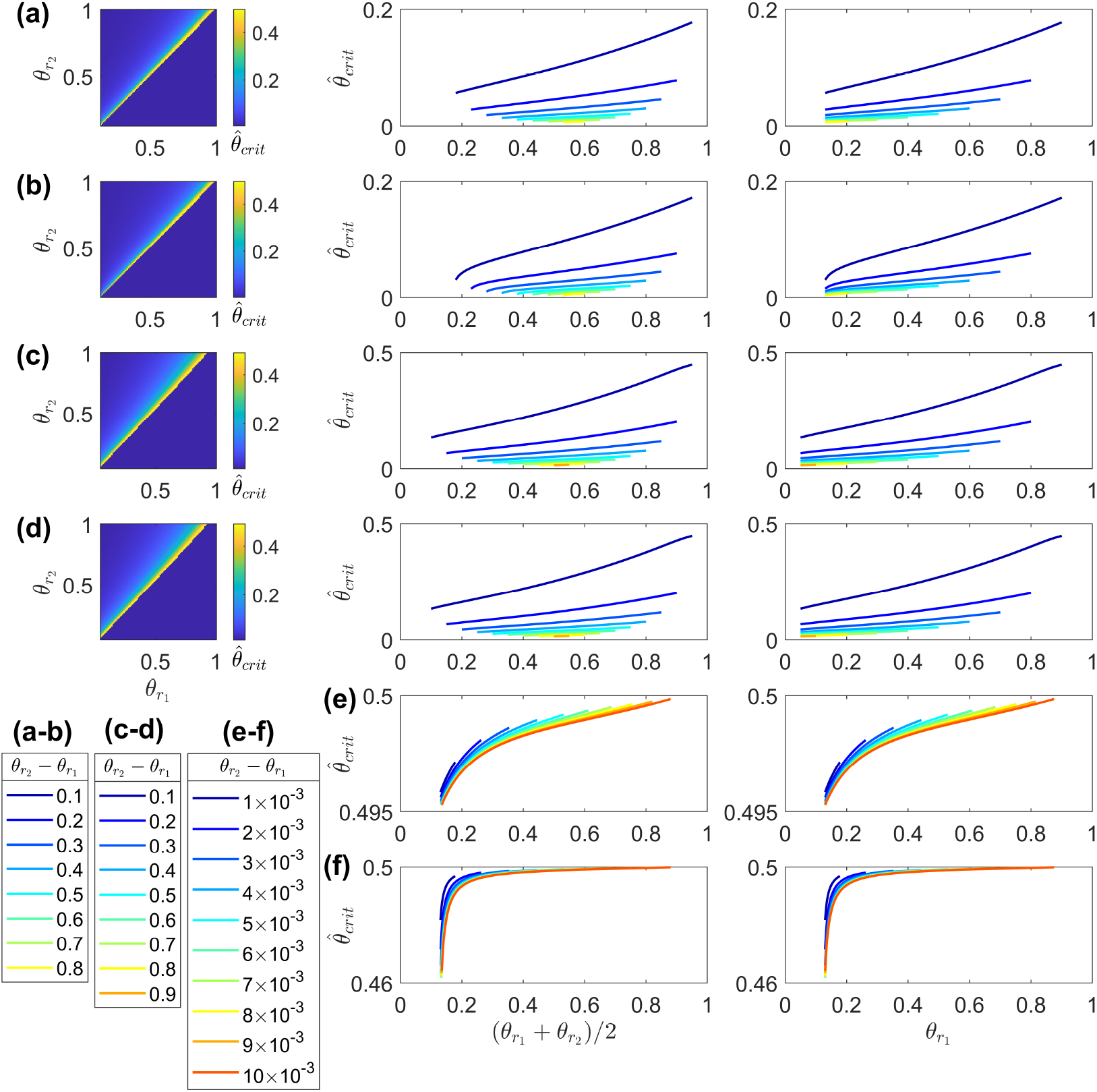
Analytical estimates of the eccentricity of the minimum trophic factor concentration using the steady-state model — wide and narrow rod degenerate patch cases. Rod loss only. All plots show the normalised eccentricity of the minimum trophic factor concentration, 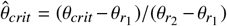. (a), (c) and (e) global treatment; (b), (d) and (f) local treatment; (a)–(d) wide patch; (e) and (f) narrow patch; (a), (b), (e) and (f) *f_crit_* = 0.3; (c) and (d) *f_crit_* = 3 × 10^−5^. (a)–(d) column 1: variation in 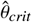 over (*θ*_*r*_1__, *θ*_*r*_2__) parameter space. (a)–(f) column 2: variation in 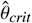 with rod degenerate patch centre position, (*θ*_*r*_1__ + *θ*_*r*_2__)/2; column 3: variation in 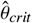 with rod degenerate patch left boundary position, *θ*_*r*_1__; each curve represents a constant rod degenerate patch width, *θ*_*r*_2__ – *θ*_*r*_1__. 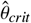 increases monotonically with decreasing rod degenerate patch width and with increasing rod degenerate patch centre/left boundary position in all cases. 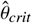 remains close to 0 for patches of width *θ*_*r*_2__ – *θ*_*r*_1__ ≥ 0.2 ((a)–(d)) and close to 0.5 for narrow patches ((e) and (f)). Analytical solutions are obtained by implicitly solving Eqs. (A.46) and (A.47) in (a) and (c), Eqs. (A.59) and (A.60) in (b) and (d), Eqs. (A.71) and (A.72) in (e), and Eqs. (A.83) and (A.84) in (f). The parameter *ϵ* = 10^−2^. Remaining parameter values as in Table 2.

## 4. Results: dynamic problem

Having explored the steady-state problem, we now consider the full dynamic problem, consisting of Eqs. (7)–(9) and (11)–(12) in the no cone regeneration/no treatment case, and Eqs. (7), (8) and (10)–(12) in the cone regeneration/treatment case. In both cases, equations are solved using the method of lines. This involves discretising in space and then integrating in time using the Matlab routine ode15s, a variable-step, variable-order solver, chosen since this is a stiff problem, involving multiple timescales. The initial trophic factor profile, *f*(*θ*, 0) = *f_init_*(*θ*), is calculated by solving Eqs. (7) and (11) at steady-state using the Matlab routine fsolve with *p_r_* = *p_r_init__*(*θ*) and *p_c_* = *p_c_init__*(*θ*).

There are a variety of scenarios to be considered:

- No rod loss patch / rod loss patch: either the retina is healthy initially or there is a single patch of rod loss;

– Wide/narrow patch: representing late and early disease stages respectively;
– Patch eccentricity: we will consider patches on the left-hand side (parafovea/perifovea), centre (mid-periphery) and right-hand side (far-periphery) of the domain;
- No mutation-induced rod loss / mutation-induced rod loss: in the absence of mutation-induced rod loss, photoreceptor (cone) degeneration is due solely to RdCVF starvation, whereas, in the presence of mutation-induced rod loss, rods also degenerate exponentially across the domain over time;
- No treatment / treatment: we consider both the natural evolution of retinal degeneration in the absence of treatment, and the effect of RdCVF treatment upon cone degeneration and cone OS regeneration;

– Global/local treatment: as in Section 3, RdCVF treatment may be applied globally, across the whole retina, or locally, within a degenerate rod patch only.

Unlike in Section 3, we do not consider the scenario in which both rods and cones have been lost from a patch initially, since, as demonstrated in that section, cone loss seldom proceeds to exceed rod loss.

We consider the following 5 cases (/combinations of scenarios):

1. Wide and narrow patches of rod loss, at left/central/right positions, without mutation-induced rod loss or treatment;
2. Mutation-induced rod loss without patch loss or treatment;
3. Wide and narrow patches of rod loss, at left/central/right positions, with mutation-induced rod loss and without treatment;
4. Wide and narrow patches of rod loss, at left-hand positions, with global and local treatment, and without mutation-induced rod loss;
5. Mutation-induced rod loss with global treatment and without patch loss.

As in Section 3, we also consider the subcases *f_crit_* = 3 × 10^−5^ and *f_crit_* = 0.3. Unlike in Section 3, we now solve and plot across the whole domain for the *f_crit_* = 0.3 subcase.

### 4.1. No treatment

Fig. 11 shows RdCVF and photoreceptor dynamics following the complete removal of rods from the interval (*θ*_*r*_1__, *θ*_*r*_2__). Cones degenerate within all wide patches of rod loss in (b) (*f_crit_* = 3 × 10^−5^) and (c) (*f_crit_* = 0.3), and within all narrow patches of rod loss in (c), but only in the left-hand narrow patch of rod loss in (b). The boundaries of cone loss remain within those of rod loss for all patches in (b) and for central and right patches in (c); however, cone loss exceeds rod loss on the left-hand side of the left wide patch and on both the left- and right-hand sides of the left narrow patch in (c). This agrees nicely with our predictions in Figs. 5 and 6.

**Figure 11:**
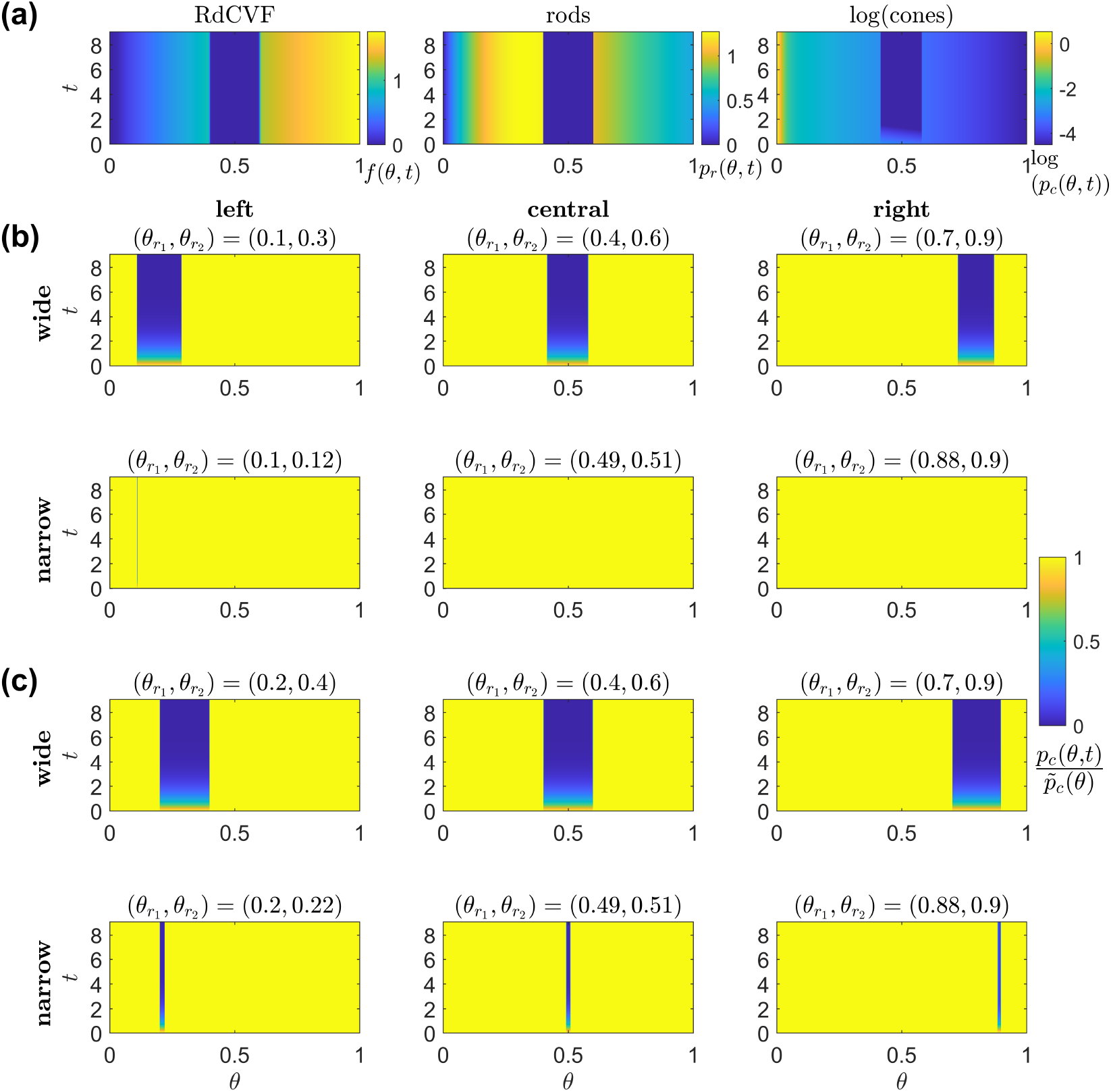
Simulations of RdCVF and photoreceptor dynamics following the complete removal of rods from the interval (*θ*_*r*_1__, *θ*_*r*_2__). (a) left-to-right: trophic factor concentration, *f* (*θ, t*), rod density, *p_r_*(*θ, t*), and the natural logarithm of the cone density, log(*p_c_*(*θ, t*)) (colour scale lower threshold: log(min(*p_c_init__*(*θ*)))), for the case *f_crit_* = 3 × 10^−5^ and (*θ*_*r*_1__, *θ*_*r*_2__) = (0.4,0.6) (the same as for the top-middle panel in (b)). (b) proportional cone loss, 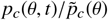, for the case *f_crit_* = 3 × 10^−5^. (c) proportional cone loss for the case *f_crit_* = 0.3. Rods are removed from wide (width *O*(1)) and narrow (width *O*(*ϵ*)) intervals, on the left-hand side (parafovea/perifovea), centre (midperiphery) and right-hand side (far-periphery) of the domain. All simulations span the period of one year in dimensional variables. Cones degenerate within all wide patches of rod loss ((b) and (c)), and within all narrow patches of rod loss in (c), but only in the left-hand narrow patch of rod loss in (b). Eqs. (7)–(9) and (11)–(12) were solved using the method of lines, with 4001 mesh points, *F*(*θ*) = *H*(*θ*_*r*_1__ – *θ*) + *H*(*θ* – *θ*_*r*_2__) and *G*(*θ*) = 1, and without treatment, cone regeneration or mutation-induced rod degeneration. The parameter = 0. Remaining parameter values as in Table 2.

Fig. 12 shows RdCVF and photoreceptor dynamics with mutation-induced rod degeneration. Cone de-generation initiates at the fovea (*θ* = 0) in (a) (*f_crit_* = 3 × 10^−5^) and at *θ* = 0.13 in (b) (*f_crit_* = 0.3), spreading peripherally (rightwards) in both cases. The retina is mostly preserved for *θ* ≤ 0.13 in (b) due to the lower value of *f_crit_* = *f*_*crit*_1__ = 3 × 10^−5^ in that region (see discussion in Section 3.2 for more details); however, given enough time, photoreceptors will also degenerate in this region, as in (a). The rate at which cone degeneration spreads across the domain increases over time as it propagates peripherally.

**Figure 12:**
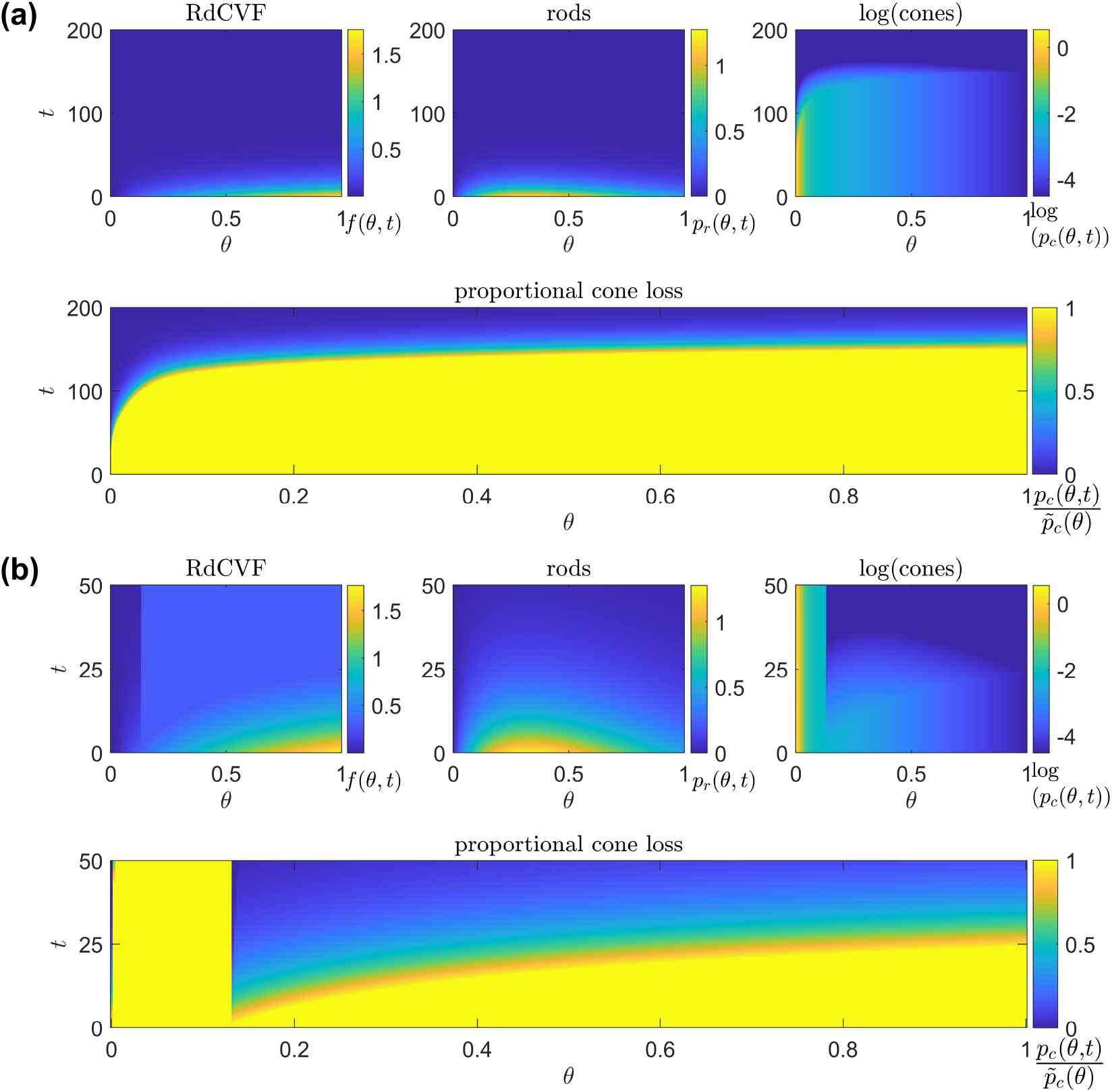
Simulations of RdCVF and photoreceptor dynamics with mutation-induced rod degeneration. (a) and (b) top, left-to-right: trophic factor concentration, *f*(*θ, t*), rod density, *p_r_* (*θ, t*), and the natural logarithm of the cone density, log(*p_c_* (*θ, t*)) (colour scale lower threshold: log(min(*p_c_init__*(*θ*)))); bottom: proportional cone loss, 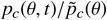. (a) *f_crit_* = 3 × 10^−5^, simulation spans ~22.1 years in dimensional variables, (b) *f_crit_* = 0.3, simulation spans ~5.5 years in dimensional variables. Cone degeneration initiates at the fovea (*θ* = 0) in (a) and at *θ* = 0.13 in (b), spreading peripherally (rightwards) in both cases. Eqs. (7)–(9) and (11)–(12) were solved using the method of lines, with *F*(*θ*) = 1 and *G*(*θ*) = 1, and without treatment or cone regeneration. (a) 4001 mesh points, (b) 401 mesh points. The parameter = 0. Remaining parameter values as in Table 2.

Fig. 13 shows RdCVF and photoreceptor dynamics with mutation-induced rod degeneration, following the complete removal of rods from the interval (*θ*_*r*_1__, *θ*_*r*_2__). Cone degeneration initiates at the fovea (*θ* = 0) in (a) (*f_crit_* = 3 × 10^−5^) and at *θ* = 0.13 in (b) (*f_crit_* = 0.3), spreading peripherally (rightwards) in both cases as in Fig. 13. Mutation-induced rod loss drives cone degenerate patch expansion and precipitates the formation of cone degenerate patches where they did not previously form ((b) centre and right narrow patches).

**Figure 13:**
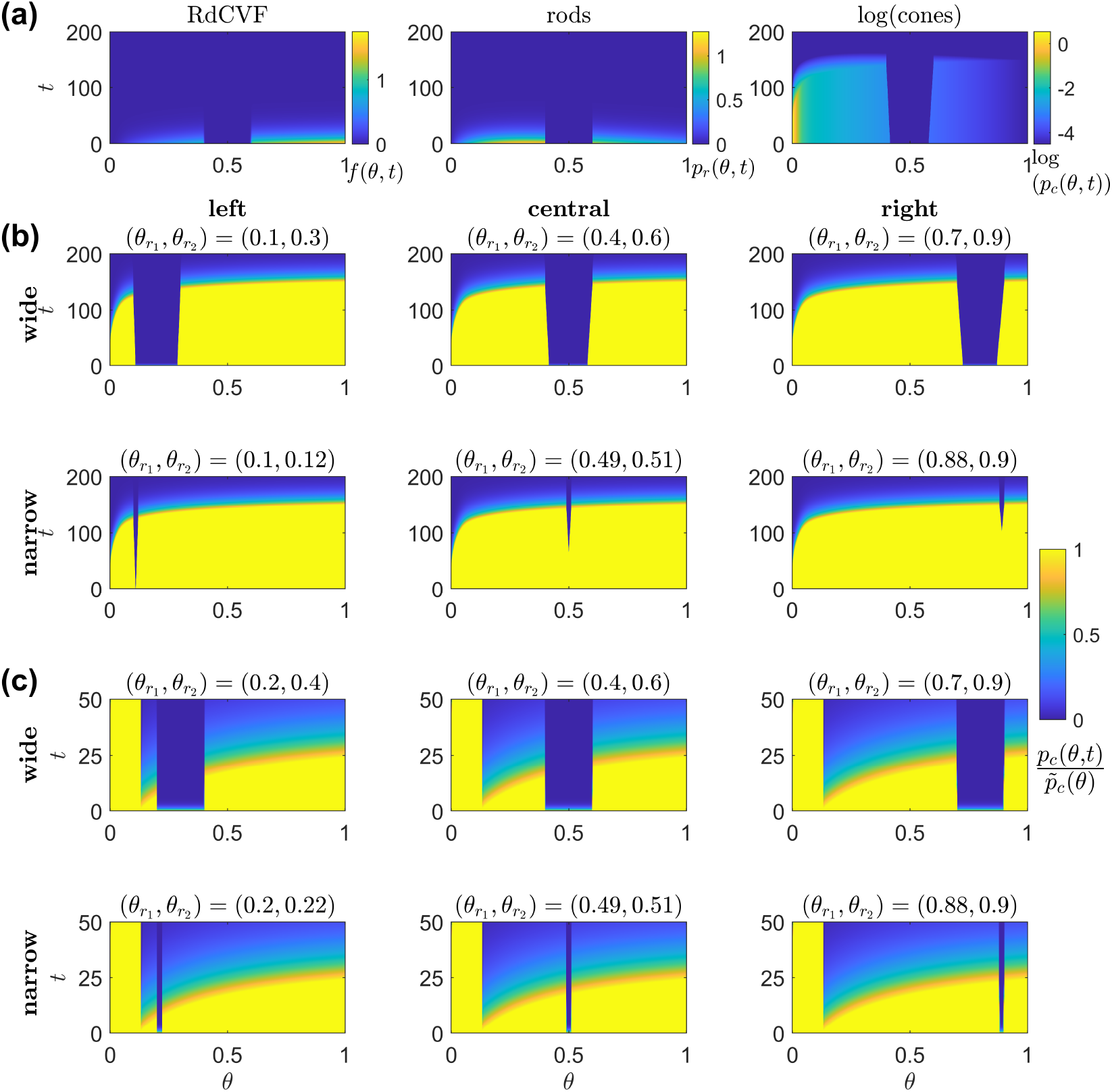
Simulations of RdCVF and photoreceptor dynamics with mutation-induced rod degeneration, following the complete removal of rods from the interval (*θ*_*r*_1__, *θ*_*r*_2__). (a) left-to-right: trophic factor concentration, *f*(*θ, t*), rod density, *p_r_*(*θ, t*), and the natural logarithm of the cone density, log(*p_c_*(*θ, t*)) (colour scale lower threshold: log(min(*p_c_init__* (*θ*)))), for the case *f_crit_* = 3 × 10^−5^ and (*θ*_*r*_1__, *θ*_*r*_2__) = (0.4,0.6) (the same as for the top-middle panel in (b)). (b) proportional cone loss, 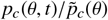, for the case *f_crit_* = 3 × 10^−5^. (c) proportional cone loss for the case *f_crit_* = 0.3. Rods are removed from wide (width *O*(1)) and narrow (width *O*(*ϵ*)) intervals, on the left-hand side (parafovea/perifovea), centre (mid-periphery) and right-hand side (far-periphery) of the domain. (a) and (b) simulation spans ~22.1 years in dimensional variables, (c) simulation spans ~5.5 years in dimensional variables. Cone degeneration initiates at the fovea (*θ* = 0) in (a) and (b), and at *θ* = 0.13 in (c), spreading peripherally (rightwards) in both cases. Patches of cone loss form and expand in all cases, initiating within patches of rod loss (compare with Fig. 11). Eqs. (7)–(9) and (11)–(12) were solved using the method of lines, with *F*(*θ*) = *H*(*θ*_*r*_1__ – *θ*) + *H*(*θ* – *θ*_*r*_2__) and *G*(*θ*) = 1, and without treatment or cone regeneration. (a) and (b) 4001 mesh points, (c) 401 mesh points. The parameter *ξ* = 0. Remaining parameter values as in Table 2.

### 4.2. Treatment

Fig. 14 shows the effects of treatment upon RdCVF and cone OS dynamics following the complete removal of rods from the interval (*θ*_*r*_1__, *θ*_*r*_2__). The treatment rate, *ξ*, is chosen to lie above the critical value, *crit*, in each case. Treatment results in the complete recovery of cone OSs in all cases, in agreement with our steady-state analytical and numerical predictions (see Figs. 7 and 8).

**Figure 14:**
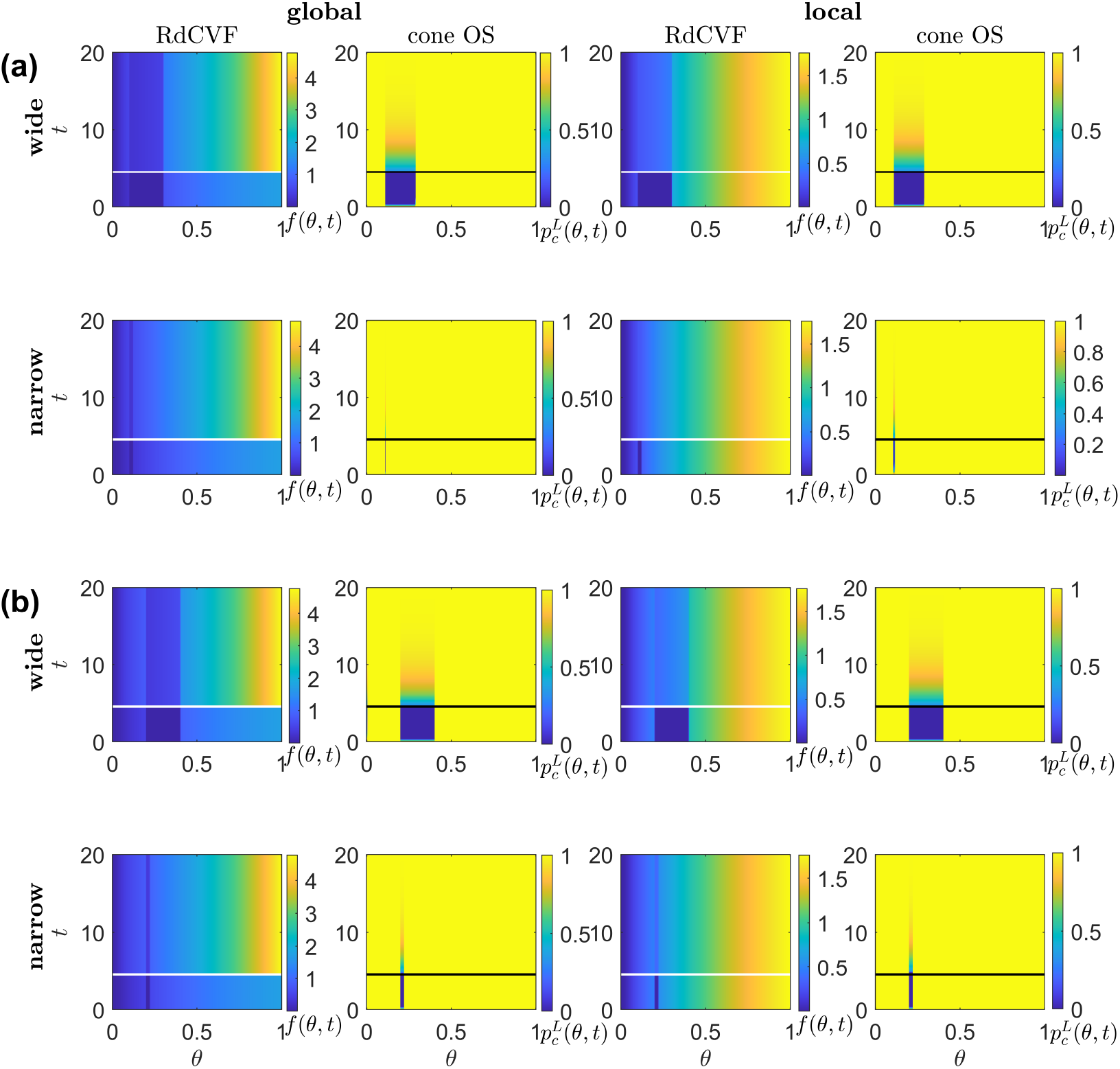
Simulations of the effects of treatment upon RdCVF and cone OS dynamics following the complete removal of rods from the interval (*θ*_*r*_1__, *θ*_*r*_2__). Panels show trophic factor concentration, *f*(*θ, t*), and cone OS length, 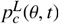, for global treatment (columns 1 and 2), *T*(*θ, t*) = *H*(*t* – *t_crit_*), and local treatment (columns 3 and 4), *T*(*θ, t*) = (1 – *F*(*θ*))*H*(*t* – *t_crit_*), following the removal of rods from wide (width *O*(1)) patches (rows 1 and 3), (a) (*θ*_*r*_1__, *θ*_*r*_2__) = (0.1,0.3) and (b) (*θ*_*r*_1__, *θ*_*r*_2__) = (0.2,0.4), and narrow (width *O*(*ϵ*)) patches (rows 2 and 4), (a) (*θ*_*r*_1__, *θ*_*r*_2__) = (0.1,0.12) and (b) (*θ*_*r*_1__, *θ*_*r*_2__) = (0.2,0.22). Black and white horizontal lines mark the time point, *t_crit_*, at which treatment is introduced. (a) *f_crit_* = 3 × 10^−5^, (b) *f_crit_* = 0.3. All simulations span the period of ~2.2 years in dimensional variables. Treatment results in the complete recovery of cone OSs in all cases. Eqs. (7), (8) and (10)–(12) were solved using the method of lines, with 401 mesh points, *F*(*θ*) = *H*(*θ*_*r*_1__ – *θ*) + *H*(*θ* – *θ*_*r*_2__) and *G*(*θ*) = 1, and without mutation-induced rod degeneration. Parameter values: *ξ* = 6 × 10^4^ and *t_crit_* = 4.53 (=1 year in dimensional variables). Remaining parameter values as in Table 2.

Fig. 15 shows the effects of treatment upon cone OS dynamics with mutation-induced rod degeneration. Strong treatment (*ξ* = 6 × 10^4^) results in the complete recovery of cone OSs in both (a) (*f_crit_* = 3 × 10^−5^) and (b) (*f_crit_* = 0.3), while weak treatment (*ξ* = 4 in (a) and = 4 × 10^4^ in (b)) delays cone OS degeneration in (a), and provides permanent cone OS recovery for *θ* ≤ 0.13 and temporary cone OS recovery for *θ* > 0.13 in (b).

**Figure 15:**
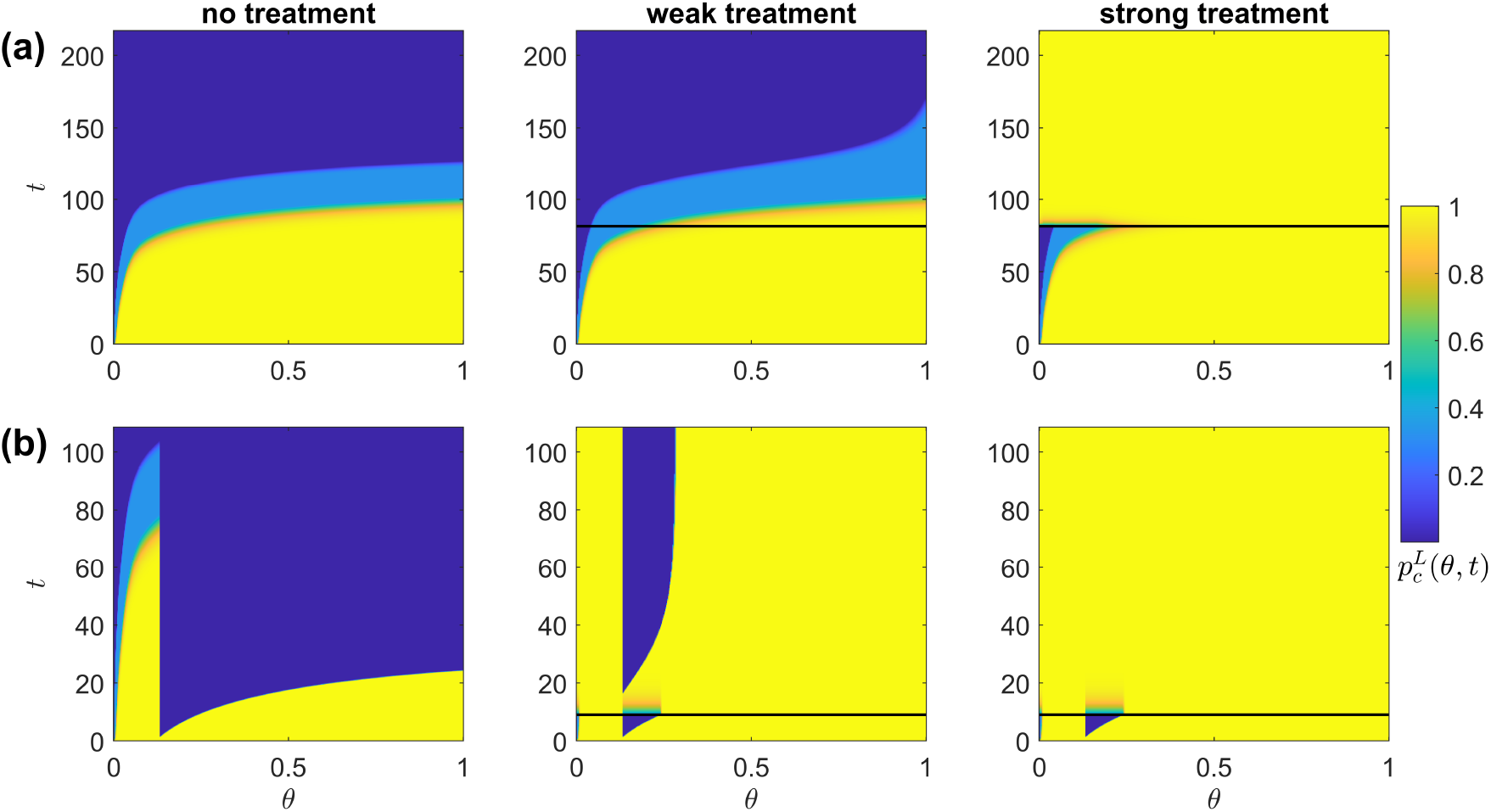
Simulations of the effects of treatment upon cone OS dynamics with mutation-induced rod degeneration. Panels show cone OS length, 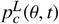, with no treatment (column 1), weak treatment (column 2) and strong treatment (column 3). Black horizontal lines mark the time point, *t_crit_*, at which treatment is introduced. (a) *f_crit_* = 3 × 10^−5^ and *t_crit_* = 81.5, while the simulation spans a period of 24 years in dimensional variables; (b) *f_crit_* = 0.3 and *t_crit_* = 9.05, while the simulation spans a period of 12 years in dimensional variables. Strong treatment results in the complete recovery of cone OSs in both (a) and (b), while weak treatment delays cone OS degeneration in (a), and provides permanent cone OS recovery for *θ* ≤ 0.13 and temporary cone OS recovery for *θ* > 0.13 in (b). Eqs. (7), (8) and (10)–(12) were solved using the method of lines, with 401 mesh points, *F*(*θ*) = 1, *G*(*θ*) = 1 and *T*(*θ, t*) = *H*(*t* – *t_crit_*). Parameter values: (a) and (b) column 1, *ξ* = 0; (a) column 2, *ξ* = 4; (b) column 2, *ξ* = 4 × 10^4^; (a) and (b) column 3, *ξ* = 6 × 10^4^. Remaining parameter values as in Table 2.

## 5. Discussion

Despite decades of research, the mechanisms underlying secondary cone loss and the spread of photoreceptor degeneration in retinitis pigmentosa (RP) have yet to be conclusively determined; though a number of candidate mechanisms have been hypothesised. One such hypothesis — the trophic factor hypothesis — suggests that rod photoreceptors produce a chemical which cone photoreceptors require to survive, such that when rods are lost, the chemical is depleted, and cone degeneration follows. Such a trophic factor, rod-derived cone viability factor (RdCVF), has been chemically identified by Léveillard et al. (2004), and was found to slow the degeneration and preserve the function of cones in rat, mouse and chick models (Fintz et al., 2003; Léveillard et al., 2004; Mohand-Saïd et al., 1998, 2000, 1997; Yang et al., 2009).

In this paper, we have formulated and analysed reaction-diffusion partial differential equation (PDE) models of RP, in which we considered RdCVF starvation as a potential mechanism of cone loss. As such, this paper complements empirical and clinical studies by isolating a biochemical mechanism in a way that would be almost impossible to achieve experimentally, allowing us to predict the *in vivo* dynamics of photoreceptor loss that would result if this mechanism (RdCVF starvation) alone were responsible for retinal degeneration (subsequent to the direct effects of the underlying mutation). We used our models to answer three key questions: (i) under what conditions will the loss of a patch of rods lead to cone degeneration?, (ii) what is the critical trophic factor treatment dose required to prevent cone degeneration following the loss of a patch of rods?, and (iii) what spatio-temporal patterns of cone degeneration can be explained via the trophic factor mechanism alone?

We began by considering the steady-state model (Section 3), first without (Section 3.2.1) and then with (Section 3.2.2) RdCVF treatment, using a combination of numerical solutions and asymptotic analysis, achieving excellent agreement between the two.

In the untreated scenario, we considered four cases: (i) wide patch of rod loss, (ii) narrow patch of rod loss, (iii) wide patch of rod and cone loss, and (iv) narrow patch of rod and cone loss; and for each of these cases, the two sub-cases: (a) low trophic factor threshold concentration, *f_crit_* = 3 × 10^−5^, and (b) high trophic factor threshold concentration, *f_crit_* = 0.3 (Figs. 4–6). In all cases it was found that cone degenerate patch width increases monotonically with increasing rod degenerate patch width and with decreasing rod degenerate patch centre eccentricity, while the difference between rod and cone degenerate patch widths decreases monotonically with rod degenerate patch centre eccentricity. Cone degenerate patch width was predicted to be less than rod degenerate patch width, except for wide and narrow rod loss patches with *f_crit_* = 0.3, where the cone degenerate patch width may exceed rod degenerate patch width close to the fovea. These results make sense in the light of the fact that the ratio of rods to cones increases monotonically with increasing eccentricity (Fig. 2(a)), such that trophic factor is scarcer toward the centre of the retina, leading to the formation of larger cone degenerate patches. It would be interesting to test these predictions experimentally by measuring the cone density local to patches of rod loss.

Trophic factor (RdCVF) treatment could be administered in a number of ways. RdCVF could be injected either into the vitreous (the structure filling the centre of the eye), or into the sub-retinal space between the photoreceptors and the retinal pigment epithelium (RPE; Yang et al., 2009). Injection into the vitreous is less invasive and would distribute RdCVF across the whole retina, while sub-retinal injections would target specific retinal areas and allow for a much higher dose to reach the cones (Yang et al., 2009). A downside to injections is that they would have to be regularly repeated throughout the lifetime of a patient, to maintain an adequate supply of RdCVF (Clérin et al., 2020). An alternative mode of administration might be to use encapsulated cell-based delivery, in which a polymer membrane capsule loaded with cells genetically modified to secrete trophic factor is surgically implanted into the vitreous, releasing trophic factor over a period of months to years (Musarella and MacDonald, 2011; Sieving et al., 2006; Tao et al., 2002). This would require less frequent interventions than with injections and provide a constant supply of RdCVF. This procedure has been tested in canines and humans, using ciliary neurotrophic factor producing RPE cells, and was found to slow photoreceptor degeneration and preserve visual acuity (Sieving et al., 2006; Tao et al., 2002). Lastly, retinal gene therapy, administered through intravitreal or sub-retinal injection of appropriately engineered adeno-associated virus vectors, could be used to genetically modify retinal cells other than the degenerating rods (e.g. RPE cells) to produce RdCVF, providing a constant, endogenous supply for the cones (Byrne et al., 2015; Clérin et al., 2020).

In the treated scenario, we considered two cases: (i) wide patch of rod loss, and (ii) narrow patch of rod loss; and for each of these cases, the two sub-cases: (a) *f_crit_* = 3 × 10^−5^, and (b) *f_crit_* = 0.3 (Figs. 7–10). In all cases the critical treatment rate, *ξ_crit_*, increases monotonically with decreasing rod degenerate patch centre and left boundary eccentricity. As above, this is to be expected from consideration of the ratio of rods to cones (Fig. 2(a)), as endogenously produced trophic factor is scarcer towards the retinal centre.

In the wide patch case, the critical treatment rate depends almost entirely upon the rod degenerate patch left boundary position, while being independent of rod degenerate patch width (Fig. 7). By contrast, in the narrow patch case, the critical treatment rate depends upon both the rod degenerate patch left and right boundary positions, and hence upon the rod degenerate patch width, the critical treatment rate increasing monotonically with increasing rod degenerate patch width (Fig. 8). The reason for this difference becomes clear from considering the normalised eccentricity of the minimum trophic factor concentration, 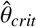, in Fig. 10. For wide patches (Fig. 10(a)–(d)), 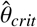 is close to the left-hand rod degenerate patch boundary, *θ*_*r*_1__, such that 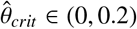 is in the left-centre-inner region (Fig. 3(a)(ii)), where only the left-hand boundary exerts an influence. Therefore, *ξ_crit_* only depends on the value of *θ*_*r*_1__ in this case. Whereas, for narrow patches (Fig. 10(e)–(f)), 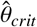 is close to the centre of the rod degenerate patch, such that 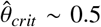 is in the centre-inner region (Fig. 3(b)(ii)), upon which both boundaries exert an influence. Therefore, *ξ_crit_* depends on the values of both *θ*_*r*_1__ and *θ*_*r*_2__ in this case.

The normalised eccentricity of the minimum trophic factor concentration, 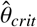, is close to the left-rather than the right-hand rod degenerate patch boundary in the wide patch case because the rod to cone ratio, and hence the trophic factor concentration, is lower there (Fig. 2(a)). 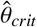 increases monotonically with decreasing rod degenerate patch width and with increasing rod degenerate patch centre and left boundary position in all cases. Knowing the location of 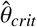 is important as it tells us which region of the retina is most at risk of degeneration (i.e. which area will degenerate first), suggesting where to spatially target treatment (e.g. with sub-retinal RdCVF injections).

Comparison of critical treatment rates in the global and local treatment cases shows that *ξ_crit_* is almost identical for global and local treatment in the wide patch case, while the critical treatment rate is higher for local than for global treatment for the narrowest patches close to the fovea in the narrow patch case (Fig. 9). Therefore, a targeted treatment (e.g. a localised sub-retinal injection of RdCVF), which uses less trophic factor, could be just as effective as a global treatment in most cases. It would be interesting to test these and the above treatment-related predictions experimentally, comparing the effects of different RdCVF doses, injection locations and local vs. global treatments.

Next we considered numerical solutions to the dynamic, time-dependent model (Section 4), first without (Section 4.1) and then with (Section 4.2) RdCVF treatment.

In the untreated scenario we considered three cases: (i) wide and narrow patches of rod loss without mutation-induced rod degeneration, (ii) mutation-induced rod degeneration without patch loss, and (iii) wide and narrow patches of rod loss with mutation-induced rod degeneration; and for each of these cases, the two sub-cases: (a) *f_crit_* = 3 × 10^−5^, and (b) *f_crit_* = 0.3 (Figs. 11–13). In Case (i), it was found that cone degeneration always occurs within wide rod degenerate patches, whereas cone degeneration may not occur in narrow rod degenerate patches away from the retinal centre for *f_crit_* = 3 × 10^−5^ (Fig. 11). Further, for *f_crit_* = 0.3, the spatial extent of cone loss may exceed that of rod loss in near-central rod degenerate patches. This is in good agreement with our predictions from the steady-state problem in Section 3.

In Case (ii), cone degeneration initiates at the retinal centre and spreads peripherally, accelerating as it does so (Fig. 12). This does not replicate any of the spatio-temporal patterns of visual field loss (and hence retinal degeneration) described by Grover et al. (1998) (see also Roberts et al., 2018). In particular, it fails to replicate the sparing of the central retina that is usually observed. This suggests that other mechanisms are necessary, either in place of or (more likely) together with trophic factor starvation-induced degeneration, to explain *in vivo* disease progression. However, we note that other spatio-temporal patterns could be generated using our model by making alternative parameter choices. For example, the rates of trophic factor production, *α*, and consumption, *β* could be reduced by two orders of magnitude, bringing the trophic factor decay term into the dominant balance with the production and consumption terms in Eqn. (7). Further, it may be that the trophic factor threshold concentration, *f_crit_*, and the rate of mutation-induced rod degeneration, *ϕ_r_*, vary with eccentricity. For example, *f_crit_* may be lower at the retinal centre if the cones in that region are protected in some way from the low RdCVF levels found there, while RdCVF supply to the central retina would be maintained for longer if *ϕ_r_* were reduced in that region (*ϕ_r_* was shown to vary with eccentricity in the healthy ageing retina by Curcio et al., 1993, thus it is reasonable to assume that it would also vary in the diseased retina, though we note that in the healthy case it is the central rods that are most vulnerable). We will explore the effects of these alternative parameter choices in a future publication.

In Case (iii), mutation-induced rod degeneration was shown to precipitate the formation of cone degenerate patches at the locations of narrow rod degenerate patches, where they would not have otherwise formed, and to drive the expansion of cone degenerate patches (Fig. 13). This occurs because the loss of rods reduces the rate of production of trophic factor, pushing RdCVF concentrations below the critical threshold in regions which were previously well-supplied.

As a hypothesis to explain the spread of photoreceptor degeneration, the trophic factor mechanism is *self-limiting*. This is because, following the loss of rods and consequent drop in RdCVF concentrations, the loss of cones decreases trophic factor demand until RdCVF levels are healthy once more. This is in contrast to the oxygen toxicity mechanism, considered by the author in previous publications (see Roberts et al., 2017, 2018), which is *self-reinforcing*; toxically high oxygen levels causing photoreceptor degeneration, leading to a further rise in oxygen levels, and so on.

In the treated scenario we considered two cases: (i) wide and narrow patches of rod loss, without mutation-induced rod degeneration, with global and local treatment, and (ii) mutation-induced rod degeneration, without patch loss, with weak and strong global treatment; and for each of these cases, the two sub-cases: (a) *f_crit_* = 3 × 10^−5^, and (b) *f_crit_* = 0.3 (Figs. 14–15). In Case (i), the treatment rate, *ξ*, was chosen to lie above the critical treatment rate, *ξ_crit_*, predicted from the steady-state model, resulting in the complete recovery of cone outer segments (OSs) in all cases as expected (Fig. 14). In Case (ii), strong treatment (*ξ* = 6 × 10^4^) resulted in the complete recovery of cone OSs, while weak treatment (*ξ* = 4 or *ξ* = 4 × 10^4^) delayed or temporarily reversed cone OS degeneration (Fig. 15). While the possible rates of RdCVF treatment have yet to be measured, our models provide useful, experimentally testable predictions of the critical treatment rate required to prevent or reverse damage to cones and the possible effects of different treatment rates.

In future work, we will extend the present study in three main directions. First, we will explore alternative scalings on the rates of RdCVF production, *α*, and consumption, *β* together with the effects of spatially heterogeneous trophic factor threshold concentration, *f_crit_*, and rate of mutation-induced rod degeneration, *ϕ_r_*, upon spatio-temporal patterns of retinal degeneration, as discussed above. Second, we will extend our models to two spatial dimensions, covering the region between the fovea and the ora serrata and incorporating variations in the azimuthal dimension (as in Roberts et al., 2018). Third, we will incorporate further biochemical details, such as those considered in Camacho et al. (2019).

In conclusion, we have constructed novel mathematical models of RP to describe and predict retinal degeneration, and the effects of treatment, under the trophic factor hypothesis. We predicted the spatial extent of cone degeneration following the loss of a patch of rods, and the critical RdCVF treatment rate required to prevent cone loss in this case. Lastly, we predicted the spatio-temporal patterns of cone degeneration that would result from this mechanism, suggesting that additional mechanisms are required to explain the patterns seen *in vivo*.

## Appendix A. Asymptotic analyses

Here we consider Cases 2-8 from Section 3.1. In each case the same approach is taken as in Section 3.1.1. Where the analysis differs from that in Section 3.1.1 we explain the difference, otherwise the results are simply stated.

In the sections that follow (Appendix A.1–Appendix A.7) we will use the following parameters (in addition to those defined in Section 3.1), defined here for ease of reference:

- 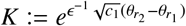;
- 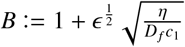;
- 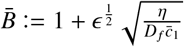;
- 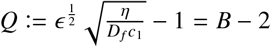;
- 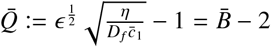.
- 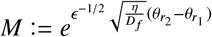.

### Appendix A.1. Narrow patch of rod loss without treatment

In this case there is no centre-outer region, and the left-centre-inner and right-centre-inner regions coalesce into a single centre-inner region (see Fig. 3(b)).

Left- and Right-Outer:

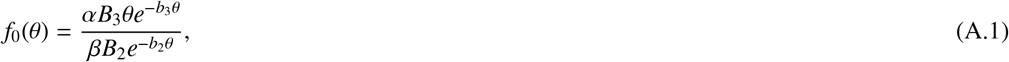

Left-Inner:

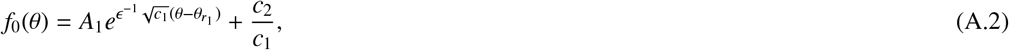

Centre-Inner:

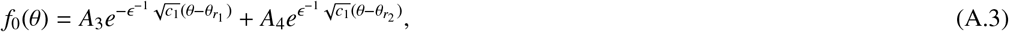

Right-Inner:

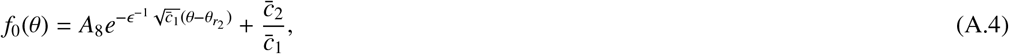

where:

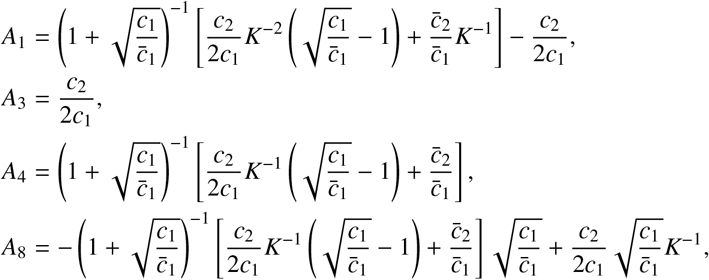

Left-Composite:

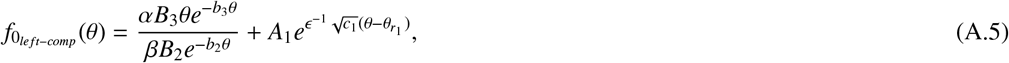

Centre-Composite is identical with the centre-inner,

Right-Composite:

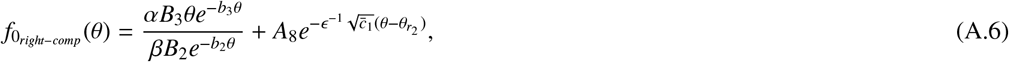

Minimal Cone Degenerate Patch Left Boundary:

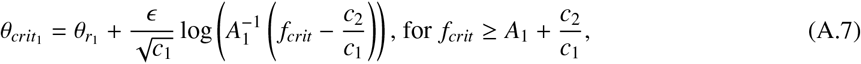

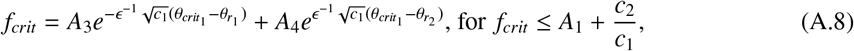

Maximal Cone Degenerate Patch Right Boundary:

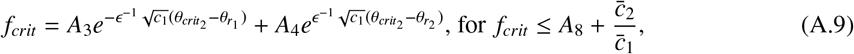

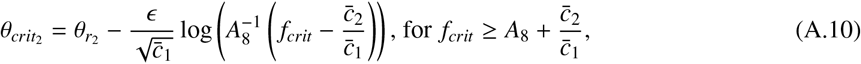

where Eqs. (A.8) and (A.9) must be solved implicitly for *θ*_*crit*_1__ and *θ*_*crit*_2__ respectively,

- when 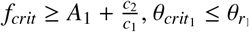;
- when 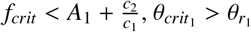;
- when 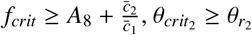;
- when 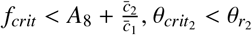.

### Appendix A.2. Wide patch of rod and cone loss without treatment

In this case, *G*(*θ*) = *H*(*θ*_*c*_1__ – *θ*) + *H*(*θ* – *θ*_*c*_2__) (and *F*(*θ*) = *H*(*θ*_*r*_1__ – *θ*) + *H*(*θ* – *θ*_*r*_2__) as before), where *θ*_*c*_1__ = *θ*_*r*_1__ and *θ*_*c*_2__ = *θ*_*r*_2__. The asymptotics for the rod and cone loss case, both here and in Appendix A.3, is valid in the region *θ* ∈ (∼0.16,1 – *ξ*).

We decompose into the same regions as in Section 3.1.1 (see Fig. 3(c)); however, the left- and rightcentre-inner regions are *O*(*ξ*^1/2^) width in this case. This is because, in the absence of cones in the central region, we must seek a dominant balance between the diffusion and decay terms. Thus, the new scaling on *θ* in the left-centre-inner region is: 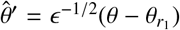; while the new scaling on *θ* in right-centre-inner region is: 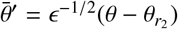; while the regular perturbation expansion for *f* in the central region is

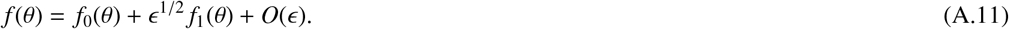

Perturbation expansions are not required for rods and cones in this region since they are absent here. The scalings on *θ* and the regular perturbation expansions in the left-inner and right-inner regions are the same as in 3.1.1.

Left- and Right-Outer:

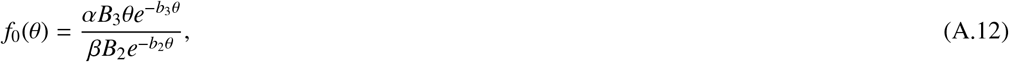

Left-Inner:

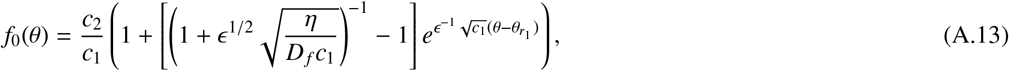

Left-Centre-Inner:

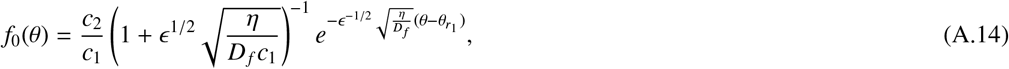

Centre-Outer:

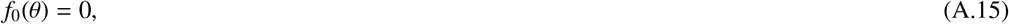

Right-Centre-Inner:

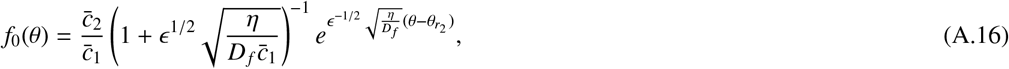

Right-Inner:

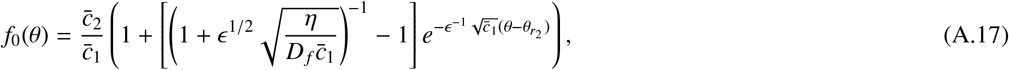

Left-Composite:

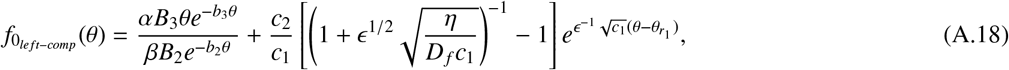

Centre-Composite:

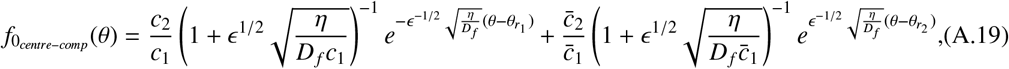

Right-Composite:

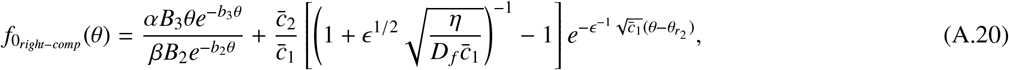

Maximal Cone Degenerate Patch Left Boundary:

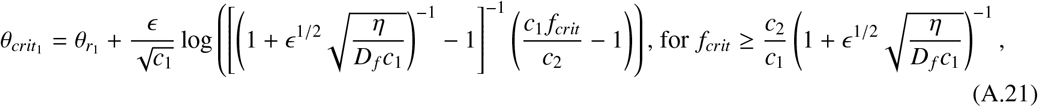

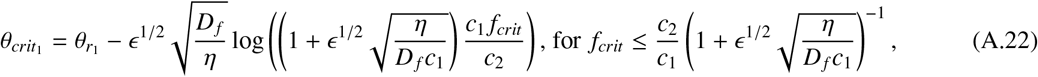

Minimal Cone Degenerate Patch Right Boundary:

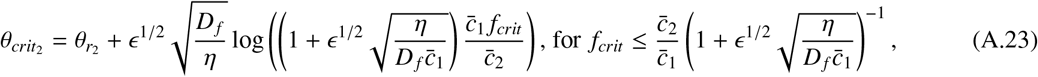

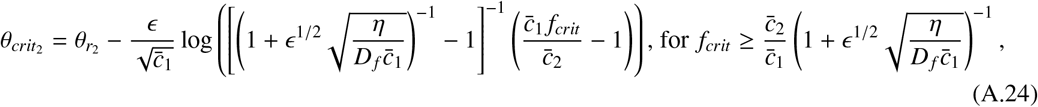

where the maximal and minimal labels above are the other way around to those in the rod loss only cases (which estimate maximal cone degenerate patch width), since here, for the rod and cone loss case, we are estimating the minimal cone degenerate patch width,

- when 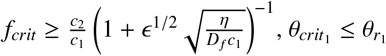;
- when 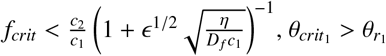;
- when 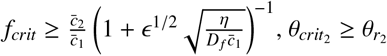;
- when 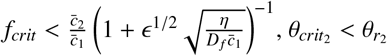.

### Appendix A.3. Narrow patch of rod and cone loss without treatment

As in Appendix A.1 there is no centre-outer region, and the left-centre-inner and right-centre-inner regions coalesce into a single centre-inner region (see Fig. 3(d)).

Left- and Right-Outer:

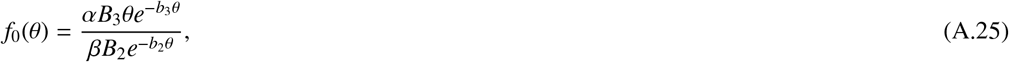

Left-Inner:

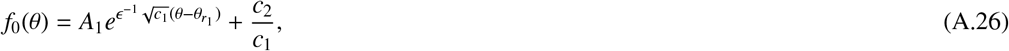

Centre-Inner:

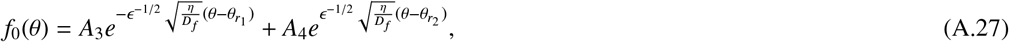

Right-Inner:

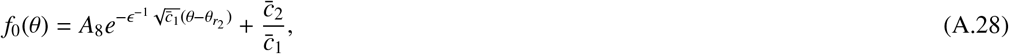

where:

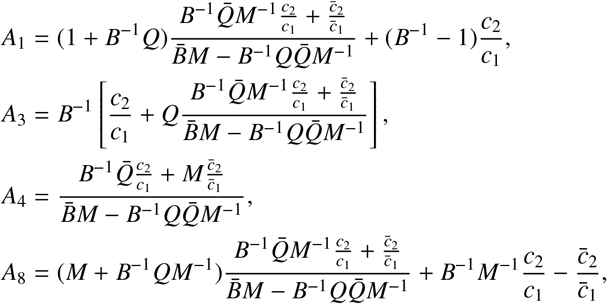

Left-Composite:

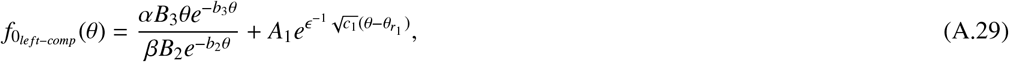

Centre-Composite is identical with the centre-inner.

Right-Composite:

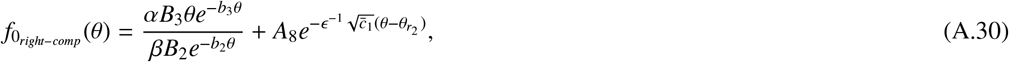

Maximal Cone Degenerate Patch Left Boundary:

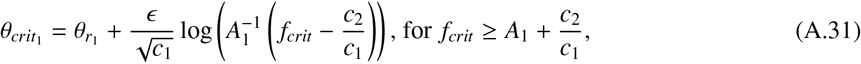

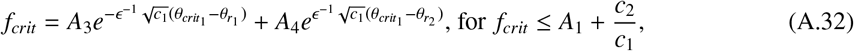

Minimal Cone Degenerate Patch Right Boundary:

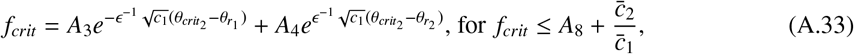

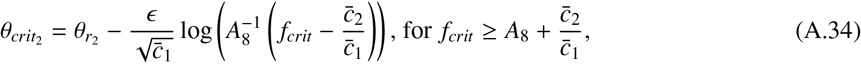

where Eqs. (A.32) and (A.33) must be solved implicitly for *θ*_*crit*_1__ and *θ*_*crit*_2__ respectively. The maximal and minimal labels above are the other way around to those in the rod loss only cases (which estimate maximal cone degenerate patch width), since here, for the rod and cone loss case, we are estimating the minimal cone degenerate patch width,

- when 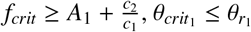;
- when 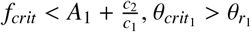;
- when 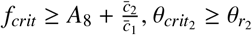;
- when 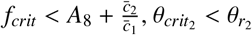.

### Appendix A.4. Wide patch of rod loss with global treatment

This case is the same as that in Section 3.1.1, except that treatment is applied across the whole domain (*T*(*θ*) = 1) at rate *ξ*. Thus, we decompose the domain into the same regions as in the untreated case (see Fig. 3(a)). We rescale *ξ* = *ϵ*^−2^ *ξ*′, where *ξ*′ = *O*(1), here and in Appendix A.5–Appendix A.7, dropping the dash.

Left- and Right-Outer:

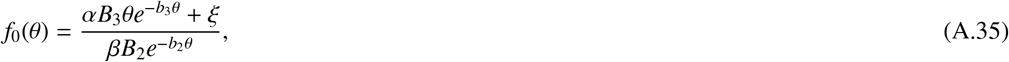

Left-Inner:

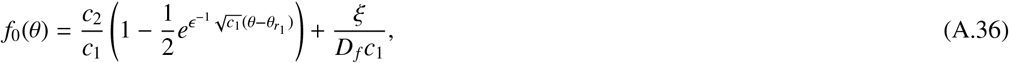

Left-Centre-Inner:

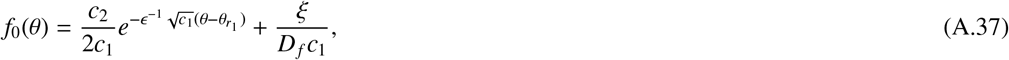

Centre-Outer:

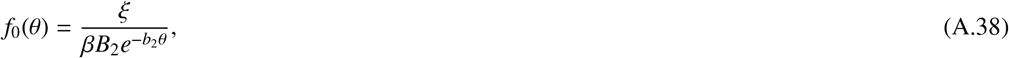

Right-Centre-Inner:

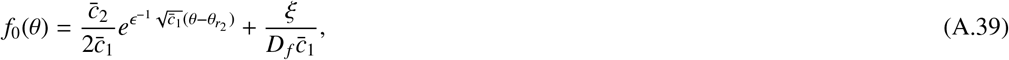

Right-Inner:

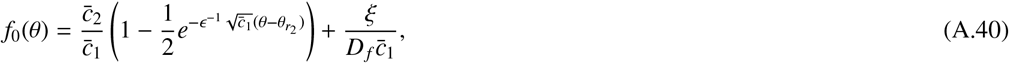

Left-Composite:

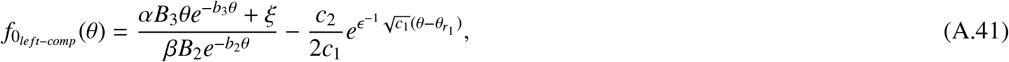

Centre-Composite:

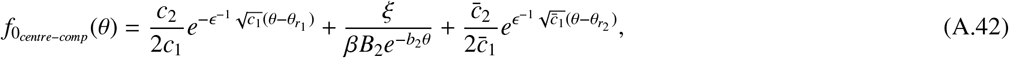

Right-Composite:

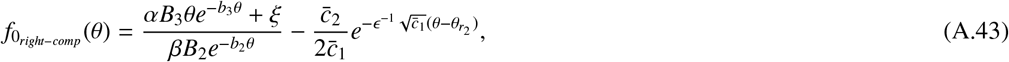

Minimal Cone Degenerate Patch Left Boundary:

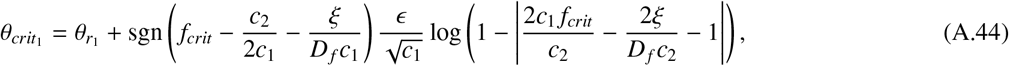

Maximal Cone Degenerate Patch Right Boundary:

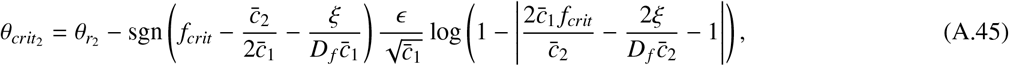

- when *f_crit_* ≥ *c*_2_/(2*c*_1_) + *ξ*/(*D_f_c*_1_), *θ*_*crit*_1__ ≤ *θ*_*r*_1__;
- when *f_crit_* ≥ *c*_2_/(2*c*_1_) + *ξ*/(*D_f_c*_1_), *θ*_*crit*_1__ > *θ*_*r*_1__;
- when 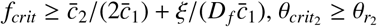;
- when 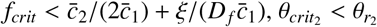.

We calculate the critical treatment rate, *ξ_crit_*, as the minimal treatment rate required to prevent cone loss, that is, to keep *f* ≥ *f_crit_* local to the patch of rod loss. The minimum trophic factor concentration is achieved at some *θ_crit_* within the interval *θ_crit_* ∈ [*θ*_*r*_1__, *θ*_*r*_2__]. Thus, *ξ_crit_* and *θ_crit_* can be found by setting *ξ* = *ξ_crit_* and *θ* = *θ_crit_* in both Eqn. (A.42) and in the equation obtained by setting 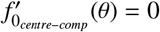 from Eqn. (A.42), to provide,

Critical treatment rate:

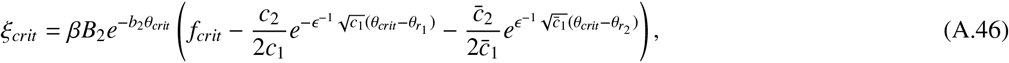

Eccentricity of the minimum trophic factor concentration:

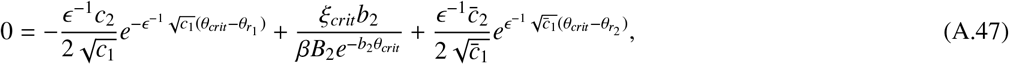

solving Eqs. (A.46) and (A.47) simultaneously.

### Appendix A.5. Wide patch of rod loss with local treatment

This case is the same as that in Appendix A.4, except that treatment is only applied locally, within the degenerate rod patch (*T*(*θ*) = 1 – *F*(*θ*)), rather than globally, across the whole domain. We decompose the domain into the same regions as in the untreated and global treatment cases (see Fig. 3(a)).

Left- and Right-Outer:

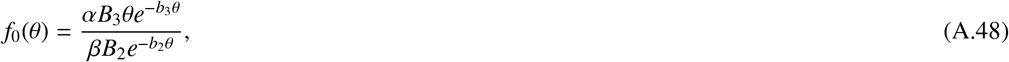

Left-Inner:

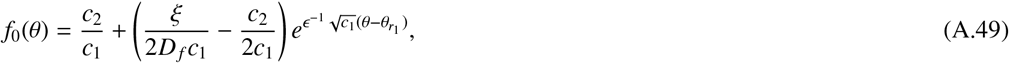

Left-Centre-Inner:

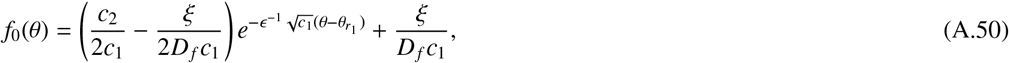

Centre-Outer:

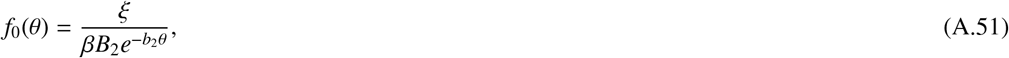

Right-Centre-Inner:

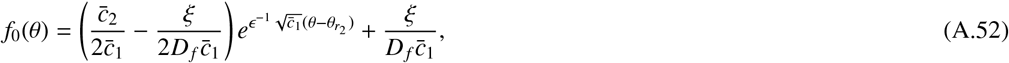

Right-Inner:

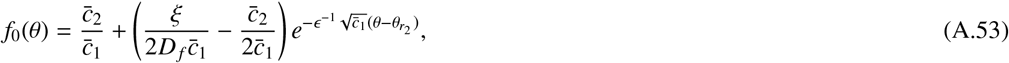

Left-Composite:

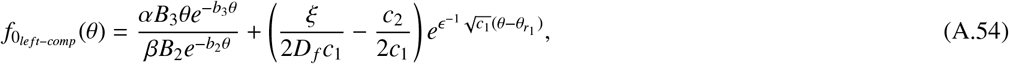

Centre-Composite:

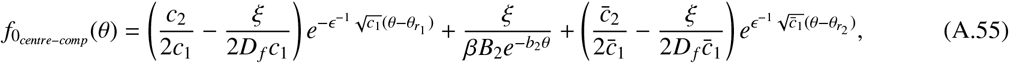

Right-Composite:

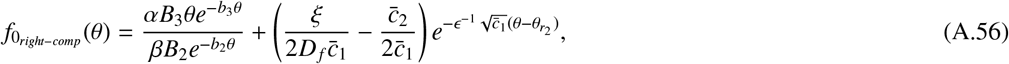

Minimal Cone Degenerate Patch Left Boundary:

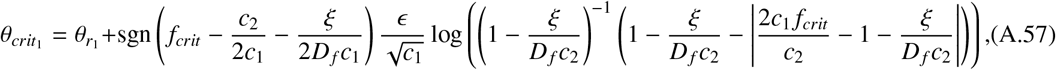

Maximal Cone Degenerate Patch Right Boundary:

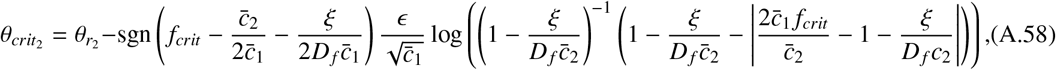

- when *f_crit_* ≥ *c*_2_/(2*c*_1_) + *ξ*/(*D_f_c*_1_), *θ*_*crit*_1__ ≤ *θ*_*r*_1__;
- when *f_crit_* ≥ *c*_2_/(2*c*_1_) + *ξ*/(*D_f_c*_1_), *θ*_*crit*_1__ > *θ*_*r*_1__;
- when 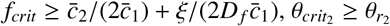;
- when 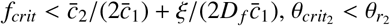.

Critical treatment rate:

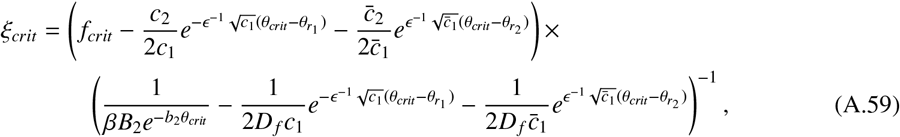

Eccentricity of the minimum trophic factor concentration:

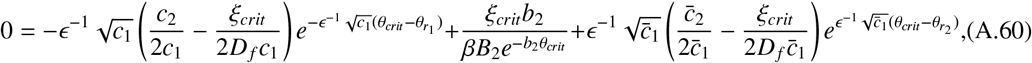

solving Eqs. (A.59) and (A.60) simultaneously for *ξ_crit_* and *θ_crit_*.

### Appendix A.6. Narrow patch of rod loss with global treatment

This case is the same as that in Appendix A.1, except that treatment is applied across the whole domain (*T*(*θ*) = 1) at rate *ξ*. Thus, we decompose the domain into the same regions as in the untreated case (see Fig. 3(b)).

Left- and Right-Outer:

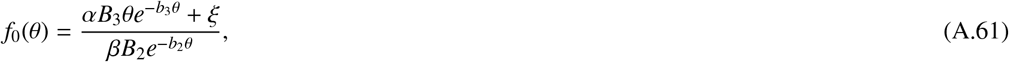

Left-Inner:

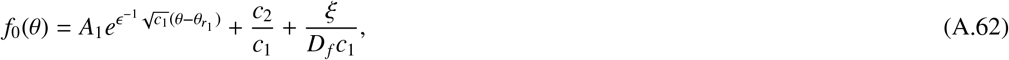

Centre-Inner:

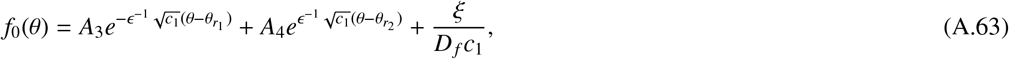

Right-Inner:

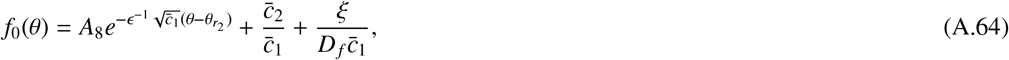

where:

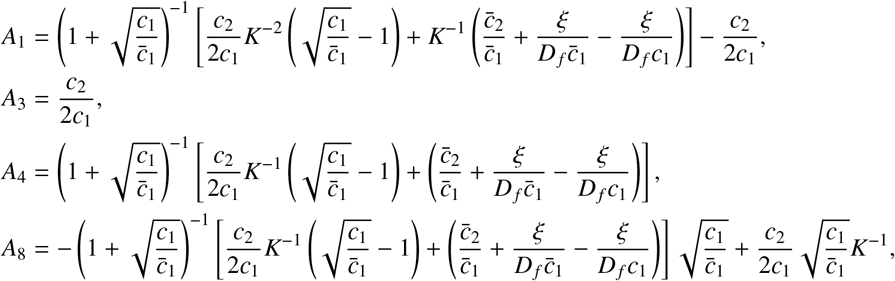

Left-Composite:

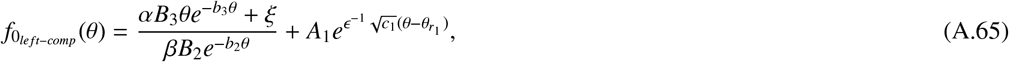

Centre-Composite is identical with the centre-inner.

Right-Composite:

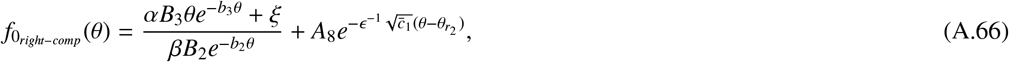

Minimal Cone Degenerate Patch Left Boundary:

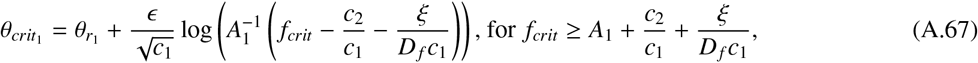

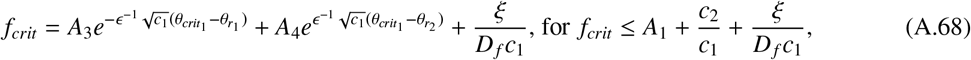

Maximal Cone Degenerate Patch Right Boundary:

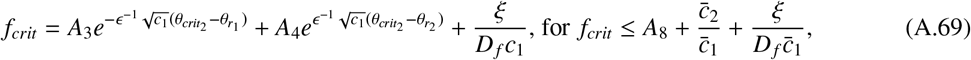

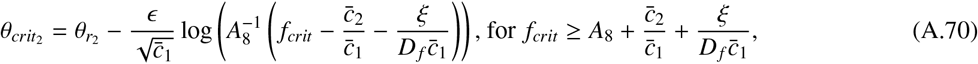

where Eqs. (A.68) and (A.69) must be solved implicitly for *θ*_*crit*_1__ and *θ*_*crit*_2__ respectively,

- when 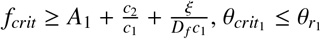;
- when 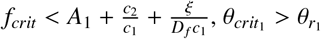;
- when 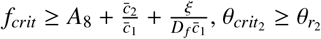;
- when 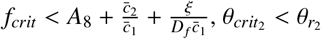.

Critical treatment rate:

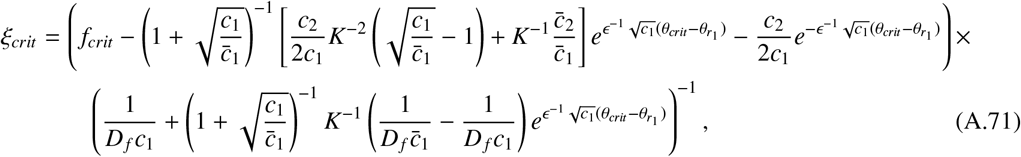

Eccentricity of the minimum trophic factor concentration:

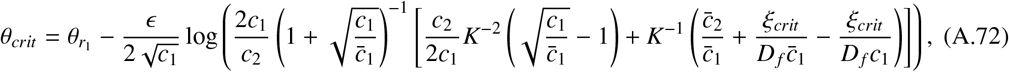

solving Eqs. (A.71) and (A.72) simultaneously for *ξ_crit_* and *θ_crit_*.

### Appendix A.7. Narrow patch of rod loss with local treatment

This case is the same as that in Appendix A.6, except that treatment is only applied locally, within the degenerate rod patch (*T*(*θ*) = 1 – *F*(*θ*)), rather than globally, across the whole domain. We decompose the domain into the same regions as in the untreated and global treatment cases (see Fig. 3(b)).

Left- and Right-Outer:

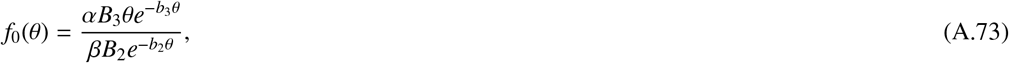

Left-Inner:

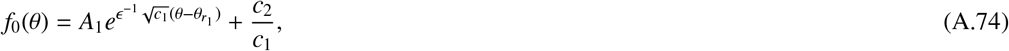

Centre-Inner:

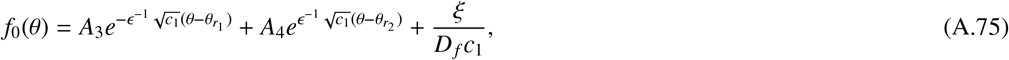

Right-Inner:

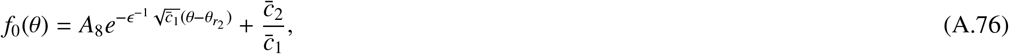

where:

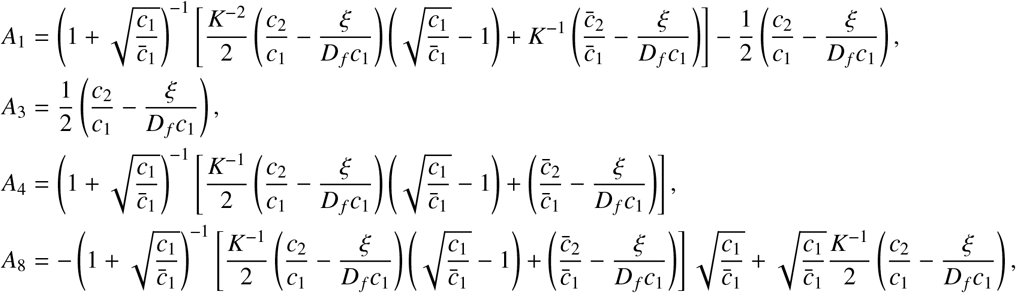

Left-Composite:

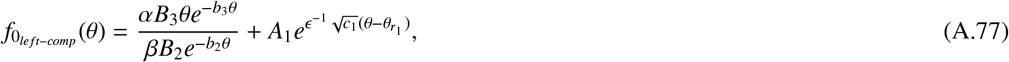

Centre-Composite is identical with the centre-inner.

Right-Composite:

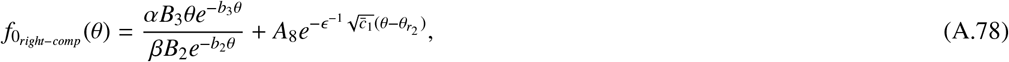

The Left- and Right-Composite solutions are the same here as for the narrow patch without treatment, except that the expressions for *A*_1_ and *A*_8_ are different.

Minimal Cone Degenerate Patch Left Boundary:

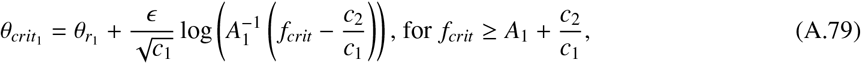

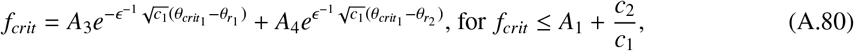

Maximal Cone Degenerate Patch Right Boundary:

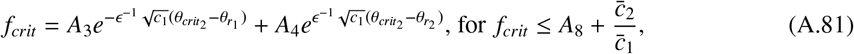

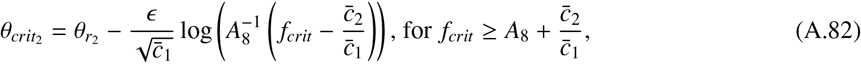

where Eqs. (A.80) and (A.81) must be solved implicitly for *θ*_*crit*_1__ and *θ*_*crit*_2__ respectively,

- when 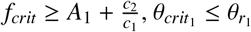;
- when 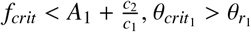;
- when 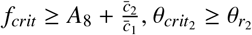;
- when 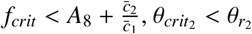.

Critical treatment rate:

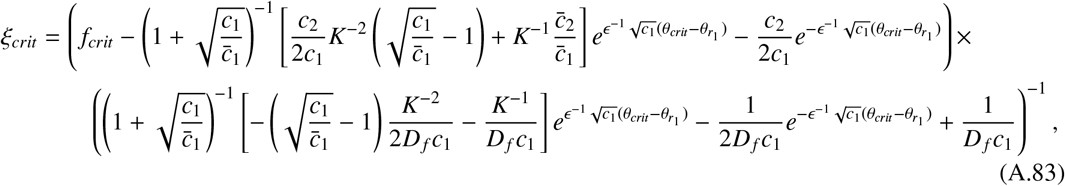

Eccentricity of the minimum trophic factor concentration:

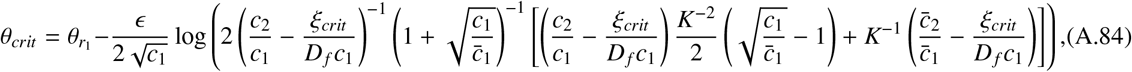

solving Eqs. (A.83) and (A.84) simultaneously for *ξ_crit_* and *θ_crit_*.

## Acknowledgements

P.A.R. is funded by BBSRC (BB/R014817/1) and thanks Tom Baden for allowing the time to pursue this research.

