## Supplementary Material for "Mathematical Models of Retinitis Pigmentosa: The Trophic Factor Hypothesis"

Paul A. Roberts*[1]

[1]School of Life Sciences, University of Sussex, John Maynard Smith Building, Brighton, BN1 9QG, UK

### S1 Justification of parameter values

- **Retinal radial position, $R$:** chosen to be $1.2 \times 10^{-2}$ m, the average radius of the human eye (Oyster, 1999).
- **Eccentricity of the ora serrata, $\Theta$:** chosen to be 1.33 rad, the extent of the retina along the horizontal meridian in the temporal direction, as measured by Curcio et al. (1990).
- **Trophic factor diffusivity, $D_f$:** the protein RdCVF has been found to come in four forms, two short forms, RdCVF-S and RdCVF2-S, and two long forms, RdCVF-L and RdCVF2-L (Chalmel et al., 2007; Léveillard et al., 2004). It is the short forms that have been found to have a role in maintaining cone viability and the first of these, RdCVF-S, which is the more effective of the two, RdCVF2-S acting in an additive (as opposed to a synergistic) fashion (Chalmel et al., 2007; Léveillard et al., 2004). Therefore, we include only the RdCVF-S form in our models. To the best of our knowledge, no measurements have been published for the diffusivity of RdCVF in any of its forms; however, the molecular weight of RdCVF-S has been measured to be 17 kDa (Léveillard et al., 2004). The protein myoglobin also has a molecular weight of 17 kDa (Zaia et al., 1992), and its diffusivity has been measured; therefore, we assume that the diffusivity of RdCVF-S is the same as that of myoglobin. Jürgens et al. (1994) have measured myoglobin diffusivity to be $1.17 \times 10^{-11}$ m$^2$s$^{-1}$ at 22°C in diaphragm muscle. Using the $Q_{10}$ rule[1] with $Q_{10} = 1.3$, gives a value of $1.73 \times 10^{-11}$ m$^2$s$^{-1}$ at 37°C (body temperature, Jürgens et al., 1994), which is the value we choose for $D_f$ (see also, McGuire and Secomb, 2001).
- **Rate of trophic factor decay, $\eta$:** to the best of our knowledge, this parameter has not been measured; however, Eden et al. (2011) have measured the half-life dynamics for a range of proteins in living human cells. In their experiments, cells were treated with the drug cisplatin, which reduces cell growth by 85%, such that the measured reduction in protein concentration is mainly due to degradation, rather than cell growth-induced dilution. Protein half-lives in the range 0.9–20.5 hr were measured, corresponding to exponential decay rates in the range $9.39 \times 10^{-6}$–$2.14 \times 10^{-4}$ s$^{-1}$. Similarly, Dörrbaum et al. (2018) measured protein half-lives in rat primary hippocampal, neuron-enriched and glia-enriched cultures ranging from 17 hr–110 days, the majority lying in the range 1–20 days, corresponding to exponential decay rates in the range $4.01 \times 10^{-7}$–$8.02 \times 10^{-6}$ s$^{-1}$. These values are smaller than (though of a similar order of magnitude to) those measured by Eden et al. (2011). We assume the values measured by Eden et al. (2011) to be more typical of the human retina since they were measured in human cells, taking the rate of trophic factor decay to be $5.13 \times 10^{-5}$ s$^{-1}$ (the mean of the rates corresponding to the half-lives measured by Eden et al., 2011).

*Corresponding author

[1] $Q_{10} = (\frac{D_2}{D_1})^{(\frac{10}{T_2 - T_1})}$, where $D_1$ ($D_2$) is the diffusivity at temperature $T_1$ ($T_2$).

- **Rate of trophic factor consumption by cones, $\beta$:** to the best of our knowledge, this parameter has not been measured. We choose the rate of trophic factor consumption such that its dimensionless value is four orders of magnitude greater than the rate of trophic factor decay, $\eta$, ensuring that the trophic factor consumption term dominates over the decay term, as would be expected biologically. Thus, we choose the dimensionless value $\beta^* = 1.79 \times 10^6$, which corresponds to a dimensional value of $\beta = 4.62 \times 10^{-12}$ m$^2$photoreceptors$^{-1}$s$^{-1}$.

- **Rate of trophic factor production by rods, $\alpha$:** to the best of our knowledge, this parameter has not been measured. In the absence of further information, we choose $\alpha$ such that the trophic factor production term is of the same order of magnitude as (and hence balances with) the consumption term, as would be expected biologically. This leaves a degree of freedom, and we choose $\alpha$ such that the mean trophic factor concentration $\tilde{f}_A = 1 \times 10^{-4}$ M, for numerical convenience. Thus, we choose $\alpha = 1.81 \times 10^{-17}$ Mm$^2$photoreceptors$^{-1}$s$^{-1}$.

- **Rate of trophic factor supply from treatment, $\xi$:** to the best of our knowledge, this parameter has not been measured. Therefore, we choose values on the order of magnitude of the critical treatment rate, $\xi_{crit}$, as predicted by numerical and analytical solutions to the steady-state problem (see Section 3.2.2).

- **Rate of mutation-induced rod degeneration, $\phi_r$:** the rate of rod degeneration in the healthy human retina has been investigated by Curcio et al. (1993), who measured a 31% reduction in the total number of rods in the central 28.5 degrees of vision between the ages of 34 and 90 yr, corresponding to an exponential decay rate of 2.10 $\times$ 10$^{-10}$ s$^{-1}$. Since the rate of rod degeneration is accelerated in RP we take this value as a lower bound and assume that the rate of mutation-induced rod degeneration is two orders of magnitude higher, that is $\phi_r = 2.10 \times 10^{-8}$ s$^{-1}$. This places the timescale of the resultant cone loss on the order of decades (see Fig. 12(a)), in line with *in vivo* progression rates.

- **Growth rate of cone OS (phases 1 and 2), $\mu_1$ and $\mu_2$:** Guérin et al. (1993) measured cone OS regrowth in rhesus monkeys following a 7-day retinal detachment period. As described in Section 2, we found that cone OS regrowth is well-described by a two-phase model. Phase 1, constant growth, occurs for cone OS lengths between 0 and 0.33 as a proportion of their full length, while phase 2, hyperbolic growth, occurs for cone OS lengths between 0.33 and 1 as a proportion of their full length (see Section 2 for more details). Cone OS length was zero immediately following reattachment (at 0 days) and reached 33% of its healthy length after 7 days. Thus, we can directly calculate the phase 1 constant growth rate as $\mu_1 = 1.60 \times 10^{-11}$ ms$^{-1}$. The phase 2 growth term was fitted to the data using the Matlab routine `fminsearch`, taking the data point at 7 days (33% length) as the initial condition and seeking to minimise the mean squared error between the model and the remaining data points to give $\mu_2 = 3.07 \times 10^{-12}$ ms$^{-1}$ (see Fig. S1). We note that it takes much longer to regrow a cone OS following the re-establishment of RdCVF supply than to completely shed it during RdCVF deprivation since in the latter case shedding occurs in the absence of regeneration while in the former case regeneration is accompanied by shedding for lengths greater than 33%, slowing the recovery of OS length (which reaches 80% of the healthy length by day 150, Guérin et al., 1993).

- **Rate of trophic factor starvation-induced cone OS degeneration, $\delta_L$:** while the rate of cone OS degeneration due specifically to RdCVF starvation has not, to the best of our knowledge, been measured, the rate of cone OS shedding under healthy conditions in humans and a number of animal species has been measured. We assume that, in the absence of sufficient RdCVF, cone OS regeneration ceases, while cone OS shedding continues at its healthy rate. Kocaoglu et al. (2016) studied cone OS shedding in the living human eye. They found that it takes between 12.67 and 16.7 days to shed (and hence renew) an entire cone OS. Based on the values given in Kocaoglu et al. (2016), we assume an average time of 14.5 days ($1.25 \times 10^6$ s) to shed a full cone OS, corresponding to a rate of $\delta_L = 2.34 \times 10^{-11}$ ms$^{-1}$. These measurements agree well with the rates of cone OS regeneration measured by Jonnal et al. (2010, 2012) and Pircher et al. (2011) in living humans, which balances shedding to maintain a roughly constant OS length.

- **Rate of trophic factor starvation-induced cone degeneration, $\delta$:** the rate of cone degeneration due specifically to RdCVF starvation has not, to the best of our knowledge, been measured; however, given that cone OS would be lost in about 14.5 days (see discussion above, Kocaoglu et al., 2016) and given that cones may be assumed to be mostly healthy immediately following OS loss, taking further time for the rest of the cell to

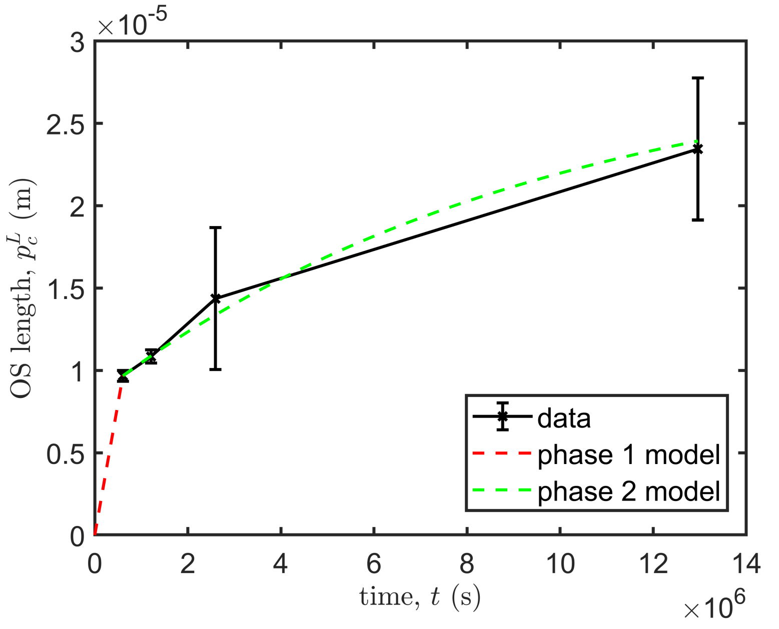


Figure S1: Graph to show model fit to cone OS regrowth data from Guérin et al. (1993). Mean data are plotted as crosses, while the error bars demarcate the standard deviation. Phase 1 — constant growth: $\dot{p}_c^L = \mu_1$; and phase 2 — hyperbolic growth: $\dot{p}_c^L = \mu_2(1 - p_c^L/\tilde{p}_c^L)$, where dot($\dot{}$) denotes the temporal partial derivative. Parameter values: $\mu_1 = 1.60 \times 10^{-11}$ ms$^{-1}$, $\mu_2 = 3.07 \times 10^{-12}$ ms$^{-1}$ and $\tilde{p}_c^L = 2.93 \times 10^{-5}$ m.

degenerate, a half-life of 28 days seems reasonable. This results in a cone degeneration rate of $2.87 \times 10^{-7}$ s$^{-1}$ (noting that half-lives from 10–100 days all result in rates of degeneration of $\sim O(10^{-7})$ s$^{-1}$).

- **Healthy cone OS length, $\tilde{p}_c^L$:** Kocaoglu et al. (2016) measured cone OS lengths in the eyes of three healthy human subjects, the average lengths being $25.5 \pm 1.7 \times 10^{-6}$ m, $30.9 \pm 1.8 \times 10^{-6}$ m and $31.6 \pm 1.5 \times 10^{-6}$ m. We take the healthy cone OS length to be the mean of these values, $\tilde{p}_c^L = 29.3 \times 10^{-6}$ m. We note that while we have chosen a biologically realistic value for this parameter for the dimensional model, the value chosen makes no difference to the parameter values in the non-dimensional model due to the way in which the parameter is cancelled out through non-dimensionalisation.

- **Trophic factor threshold concentration, $f_{crit}$:** the minimal RdCVF concentration required to maintain cones in health has not, to the best of our knowledge, been measured. In the absence of further information, we choose two possible values for $f_{crit}$. The first value ($f_{crit} = 3 \times 10^{-9}$ M) is taken to lie just below the minimum RdCVF concentration at steady-state under healthy conditions and in the absence of treatment (i.e. where $p_r = \tilde{p}_r(\theta)$, $p_c = \tilde{p}_c(\theta)$ and $\xi = 0$), which occurs at the centre of the fovea ($\theta = 0$ rad). The second value ($f_{crit} = 3 \times 10^{-5}$ M) is taken to be $10^4$ times larger than the first value, lying just below the minimum trophic factor concentration away from the fovea ($\theta > 0.13 \times \Theta$ rad). See Section 3.2 for more details.

- **Rod and cone profile parameters, $B_1$, $B_2$, $B_3$, $b_1$, $b_2$, $b_3$:** determined by fitting the functions $p_r = \tilde{p}_r(\theta)$ and $p_c = \tilde{p}_c(\theta)$ to Curcio et al. (1990)'s measurements of the mean rod and cone distributions along the temporal horizontal meridian in healthy humans using the Trust-Region Reflective algorithm in Matlab's curve fitting toolbox.

- **Mean trophic factor concentration, $\tilde{f}_A$:** calculated as the mean RdCVF concentration in the dimensional model at steady-state under healthy conditions and in the absence of treatment (i.e. where $p_r = \tilde{p}_r(\theta)$, $p_c = \tilde{p}_c(\theta)$ and $\xi = 0$).

- **Mean photoreceptor density, $\tilde{p}_A$:** calculated as the mean photoreceptor density across the retina under healthy conditions, that is, the mean of $\tilde{p}_r(\theta) + \tilde{p}_c(\theta)$.
- **Degenerate patch boundaries (rods and cones), $\theta_{r_1}$, $\theta_{r_2}$, $\theta_{c_1}$ and $\theta_{c_2}$:** chosen to explore disease progression following the loss of patches of rods and/or cones and selected such that $0 \leq \theta_{r_1} < \theta_{r_2} \leq \Theta$ (rad) and $0 \leq \theta_{c_1} < \theta_{c_2} \leq \Theta$ (rad).
- **Position of left- and right-hand limits of local treatment, $\theta_{treat_1}$ and $\theta_{treat_2}$:** chosen to explore the effects of local trophic factor treatment upon cone degeneration and cone OS recovery. Selected such that $0 \leq \theta_{treat_1} < \theta_{treat_2} \leq \Theta$ (rad).
- **Time at which treatment first applied, $t_{crit}$:** chosen to explore the effect of trophic factor treatment at a given stage of retinal degeneration. Selected such that $t_{crit} > 0$ s.
